# Implicit Laplacian of Enhanced Edge: An Unguided Algorithm for Accurate and Automated Quantitative Analysis of Cytoskeletal Images

**DOI:** 10.1101/2021.05.11.442512

**Authors:** Pai Li, Ze Zhang, Yiying Tong, Bardees M. Foda, Brad Day

## Abstract

The eukaryotic cytoskeleton plays essential roles in cell signaling and trafficking, which is broadly associated with immunity and diseases of human and plants. To date, most analyses aiming at defining the temporal and spatial dynamics of the cytoskeleton have relied on qualitative and quantitative analysis of fluorescence images to describe cytoskeletal function. While state-of-the-art, these approaches have limitations: the diverse shape and brightness of the cytoskeleton cause considerable inaccuracy in both human-driven and automated approaches, and the widely adopted image projection process (3D to 2D) leads to bias and information loss. Here, we describe the development and application of Implicit Laplacian of Enhanced Edge (ILEE), an unguided approach that uses a 2D/3D-compatible local thresholding algorithm for the quantitative evaluation of cytoskeletal status and organization at high performance. Using ILEE, we constructed a Python library to enable automated cytoskeletal image analysis, providing numerous biologically-interpretable indices measuring the density, bundling, severing, branching, and directionality of the cytoskeleton. The data presented herein demonstrate that ILEE resolves the defects of classic cytoskeleton analysis approaches, enables the measurement of novel cytoskeletal features, and yields quantitatively descriptive data with superior accuracy, stability, and robustness. We released the ILEE algorithm as an open-source library and further developed a Google Colab interface as a community resource.

## Introduction

Higher eukaryotes have evolved a complex suite of cellular signaling mechanisms to regulate many biological processes, such as growth, development, movement, reproduction, and response to environmental stimuli. Not surprisingly, to maintain the integration and sustainability of a conceptual signaling network, another “physical network” is deployed as the framework to organize all subcellular structures and the conduit to transport molecules and information^1^ – namely, the cytoskeleton. Shared by plants and animals are two types of cytoskeleton, actin and microtubule, which can be generalized as a cytoplasmic web-like matrix, physically integrating the plasma membrane, vesicles, and organelles and functionally connecting the intercellular signaling to the extracellular environments through a highly dynamic series of temporally- and spatially-regulated changes of architecture^2,3^. Altogether, the cytoskeleton controls numerous cellular processes such as movement, shaping, cellular trafficking, and intercellular communication^4^. It also provides the mechanical force required for chromosome separation and plasma membrane division during mitosis and meiosis^5^. Besides its role within the cytoplasm, the cytoskeleton also participates in a variety of processes within the nucleus, including RNA polymerase recruitment, transcription initiation, and chromosome scaffolding^6^.

As the cytoskeleton is vital for cell activities and life, the dysfunction of cytoskeletal dynamics generally leads to severe disease. For example, in the development of Alzheimer’s disease, amyloid development triggers the dysregulation of actin dynamics within dendritic spines, leading to synaptotoxicity^7^. Similarly, the dysfunction of microtubule dynamics can trigger neuropsychiatric disorders^8^. In breast cancer, the cancer cell can deploy aberrant actin aggregation to resist cytotoxic natural killer (NK) cells from the immune system^9^. In the case of plant diseases, the function of the cytoskeleton is similarly required: recent work has demonstrated that the dynamics of actin and microtubule of the host are specifically manipulated by pathogens to paralyze the plant immunity during the infection^10^. Indeed, the eukaryotic cytoskeleton not only serves as an integrated cell signaling and transportation platform, but also take risks as “a focus of disease development” at the molecular level for human and plants. Hence, understanding the architecture and dynamics of the cytoskeleton is broadly associated with life, health, and food security, attracting significant interests across different fields of biology and beyond.

Over the past several decades, confocal microscopy-based methods using fluorescence markers have been developed to monitor changes in cytoskeletal organization^11^. While showing advantages in real-time observation and intuitive visual presentation, these approaches possess critical limitations – namely, they are subject to human interpretation, and as a result, often suffer from bias. As a step to reduce such limitation(s), the emergence of computational, algorithm-based analyses offers a solution to describe the quantitative features of cytoskeletal architecture with reduced human bias. However, while early studies introduced the concept of using generalizable image processing pipelines^12,13^ to transfer the task of evaluation away from the user and into a series of computer-based quantitative indices, several key bottlenecks were still prevalent. First, most of the quantitative algorithms that have been described to date are limited to 2D images. As a result, these approaches require users to manually generate z-axis projections from raw data, a process that results in an incredible amount of information loss, especially within the perpendicular portion of the cytoskeleton. Second, many current approaches require users to set thresholds manually to segment cytoskeletal components from the images, a task that results in sampling bias. Lastly, the performance of existing algorithms varies significantly depending on the sample source. This hurdle imposes a considerable disparity in the algorithm performance for plants – which possess a dominance of curvy and fluctuating filaments – compared to the animal cytoskeletal organization, which is generally straight and complanate^14–16^. In fact, while sample source dramatically impacts our ability to evaluate the features of cytoskeletal function across all eukaryotes, the vast majority of current approaches are developed based on cytoskeletal images from animal cells, which indicates potential systemic bias when applied to other types of image samples, such as plants.

Previous work has described the development of a global-thresholding-based pipeline to define and evaluate two key features of cytoskeleton filament organization in living plant cells: cytoskeletal density, defined by *occupancy*, and bundling, defined by statistical *skewness* of fluorescence^17^. While this approach utilizes manual global thresholding (MGT), which can potentially introduce a certain level of user bias, it still outperforms most standardized adaptive/automatic global or local thresholding approaches, such as Otsu^18^ and Niblack^19^. More recent advances in MGT-based approaches, such as those by Higaki and colleagues, include additional analysis indices, such as the introduction of coefficient of variation (*CV*) of fluorescence to quantify the degree of filament bundling – an improvement that advances the overall robustness and utility of the original algorithm^20^. However, this analysis pipeline still consumes a considerable amount of time and effort from users for massive sample processing and leaves unaddressed two critical issues of rigor in image processing and analysis: information loss and human bias.

In the current study, we developed implicit Laplacian of enhanced edge (ILEE), a 2D/3D compatible unguided local thresholding algorithm for cytoskeletal segmentation and analysis, which is based on the native brightness, first-order derivative (i.e., gradient), and second-order derivative (i.e., Laplacian) of the cytoskeleton image altogether (see Fig. 1). The study described herein supports ILEE as a superior image quantitative analytic platform that overcomes current limitations such as information loss through dimensional reduction, human bias, and inter-sample instability. As shown, ILEE can accurately recognize cytoskeleton from images with a high variation of filament brightness and thickness, such as live plant samples.

**Fig. 1:**
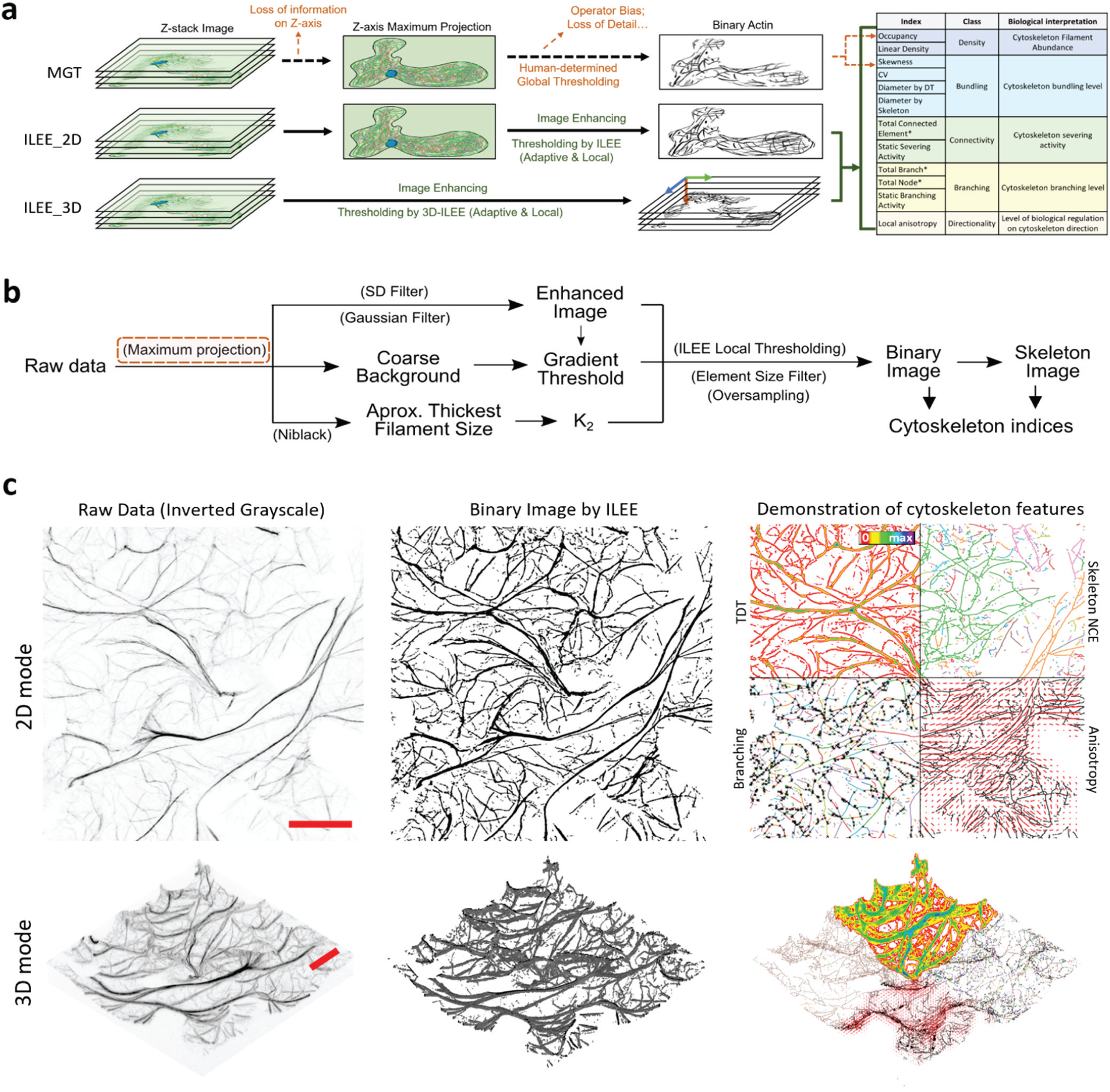
ILEE pipeline and the demonstration of cytoskeletal indices. **a**, ILEE is an adaptive local thresholding approach that applies to both 2D and 3D data structures, with an output of 12 cytoskeletal indices. Please note that *total connected element, total branch*, and *total node* (marked “*”) are not normalized indices for direct biological interpretation. **b**, Schematic diagram of the ILEE algorithm. ILEE requires a sample image with enhanced edge gradients, a computed gradient threshold, and an implicit Laplacian smoothing coefficient, *K*_*2*_, to generate a binary image and skeletonized image for index computation. Z-axis maximum projection (red box) is only conducted in the 2D mode. **c**, Visualized demonstration of ILEE performance. The raw data, binary image generated by ILEE, and demonstration of computed features by both 2D and 3D modes are shown. TDT, total distance transformation map, used to compute all diameter indices; skeleton NCE, non-connected elements of skeleton image with each element by different colors, used to estimate severing activity; branching, skeleton image with each branch in different color and each node by a black cross; anisotropy, local anisotropy level, shown as length of red lines, and direction of the first eigenvector, shown as the direction of red lines. Scale bar = 20μm.

As a key advance in the development of ILEE, we constructed an ILEE-based Python library, namely ILEE_CSK, for the fully automated quantitative analysis of cytoskeleton images. In order to improve ILEE_CSK’s capability to explore the features of cytoskeleton, we proposed several novel indices – *linear density, diameter_TDT, diameter_STD, static severing activity*, and *static branching activity* to enable/enhance the measurement of (de-)/polymerization, bundling, severing, and branching dynamics of the cytoskeleton. Together with other classic indices transplanted from other studies, ILEE_CSK totally supports 12 cytoskeletal indices within 5 primary classes: density, bundling, connectivity, branching, and directionality. Our data suggested that ILEE outcompetes other classic algorithms by its superior accuracy, stability, and robustness over the computation of cytoskeleton indices. Furthermore, we provided evidence demonstrating higher fidelity and reliability of 3D-based cytoskeletal computational approaches over traditional 2D-based approaches. In addition, using a series of experiment-based images from pathogen-infected plant cells, we demonstrated that ILEE has an improved sensitivity to distinguish biological differences and owns the potential to reveal novel biological features previously omitted due to insufficient approaches. In sum, this platform not only enables the efficient acquisition and evaluation of key actin filament parameters with high accuracy from both projected 2D and native 3D images, but also enables accessibility to a broader range of cytoskeletal status for biological interpretation.

The library, ILEE_CSK, is publicly released at GitHub (https://phylars.github.io/ILEE_CSK/). We also developed the ILEE Google Colab pipelines for data processing, visualization, and statistical analysis, which is a convenient and user-friendly interface that requires no programing experience or particular computational device.

## Results

### The ILEE pipeline

Raw images generated by laser scanning confocal microscopy are typically obtained through detecting in-focus photons by a sensor from each resolution unit on a given focal plane. Since the cytoskeleton is a 3D structure that permeates throughout the cell, current approaches used to capture the filament architecture rely on scanning the pixels of each plane along the z-axis at a given depth of step; finally, these stacks are reconstructed to yield a 3D image. However, due to limited computational biological resources, most studies have exclusively employed the z-axis-projected 2D image, which results in substantial information loss and systemic bias in downstream analyses.

In our newly developed algorithm, we integrated both 2D and 3D data structures into the same processing pipeline to ameliorate the aforementioned conflict (Fig. 1a). In short, this pipeline enabled automatic processing and evaluation of both traditional 2D and native 3D z-stack image analysis. As shown in Fig. 1b, cytoskeleton segmentation using ILEE requires three inputs: an edge-enhanced image, a global gradient threshold that recognizes the edges of potential cytoskeletal components, and the Laplacian smoothing coefficient *K* (described below). With these inputs, a local threshold image is generated via ILEE, and the pixels/voxels with values above the threshold image at the same coordinates are classified as cytoskeletal components. The output of this is the generation of the binary image (Fig. 1c). Once acquired, the binary image is further skeletonized^21^ to enable the downstream calculation of numerous cytoskeleton indices, the sum of which comprises the quantitative features of cytoskeletal dynamics (Fig. 1c). Additionally, because the 2D and 3D modes share a common workflow, all of the calculated cytoskeleton indices also share the definition for both modes, regardless of the difference in dimensional spaces. This additional feature enables a horizontal comparison of both modes by the user, which we assert will significantly contribute to the community by providing massive image datasets for further examination and comparison through the open-source library. In general, the ultimate goal of this approach, and resultant algorithm, is to construct a pipeline that enables the automated detection of the eukaryotic cytoskeleton from complex biological images in an accurate and unbiased manner.

### Image decomposition and processing strategy

One of the central problems of automated cytoskeletal image processing is how to accurately recognize cytoskeletal components – a task that is highly challenging because object pixels (i.e., cytoskeleton components) generally have a high dynamic range of signal intensity within and among individual samples, due to varied bundle thickness, concentration of fluorescent dye, and its binding efficiency. As a framework to further understand this challenge, an image from confocal microscopy is conceptually comprised of three components: (1) the true fluorescence, that which is emitted by the dye molecules within the pixel, (2) the diffraction signal transmitted from neighboring space, and (3) the ground noise generated by the sensor of the imaging system (Fig. 2a). During confocal imaging, the ground noise fluctuates around a constant due to the fixed setting of photon sensors, while the diffraction signal is positively correlated with the local true fluorescence. Therefore, an ideal cytoskeleton segregation algorithm will be an adaptive local thresholding approach that refers to both ground noise and local signal intensity.

**Fig. 2:**
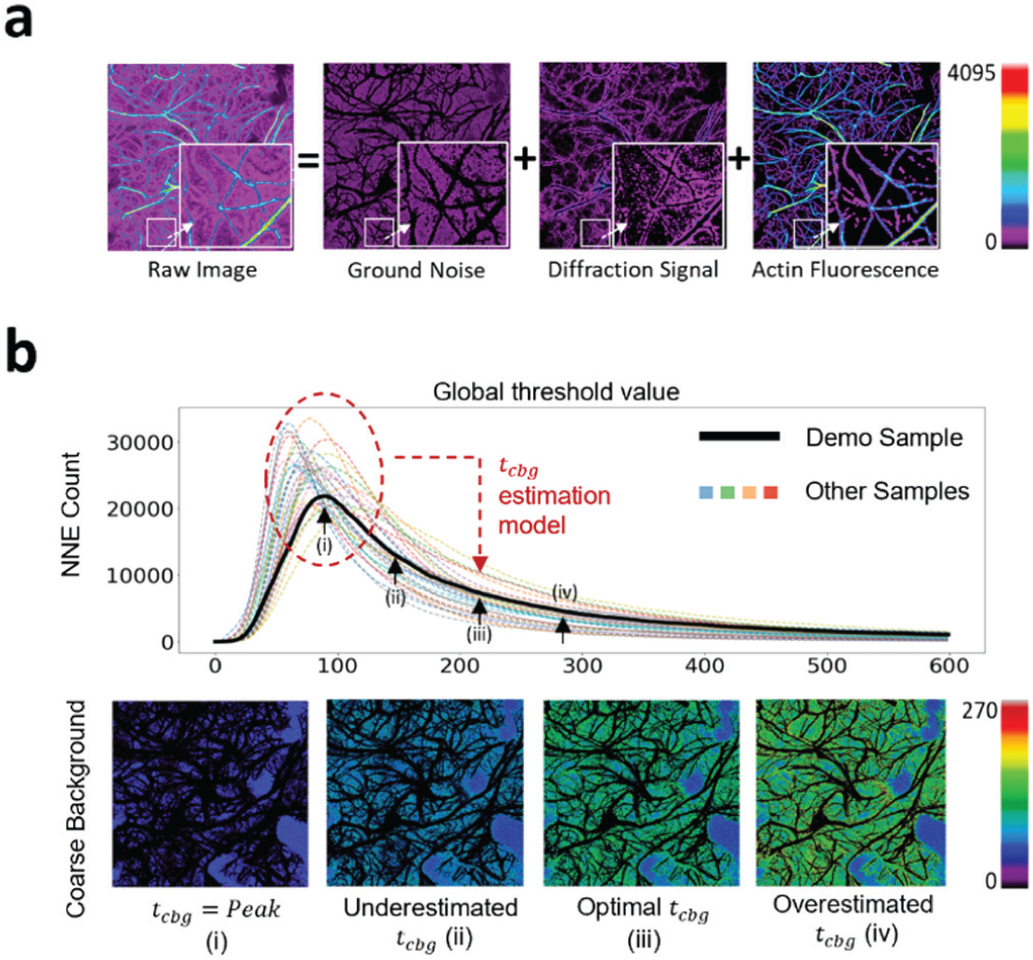
NNES global thresholding. **a**, The conceptual decomposition of a confocal fluorescence image of the cytoskeleton. An Arabidopsis leaf confocal microscopic image of actin, as an example of the eukaryotic cytoskeleton, comprises of 3 components: ground noise, the mechanical noise of the sensor regardless of the true fluorescence signal, diffraction light, the unavoidable diffraction signal of fluorescence component around, and true actin signal. They correspond to the noise filtered by coarse background, noise additionally filtered by ILEE, and segmented actin components in the algorithm. **b**, The scheme and demonstration of NNES. The curve reflects the NNE (negative non-connected component) count when certain global thresholding is applied to the raw images of 30 randomly selected samples. Their extremely smooth shape makes it easy to detect the peak as a feature value as the input for the coarse background estimation model. The demonstration of the filtered background at positions (i), (ii), (iii), and (iv) are shown above, where (iii) is adopted. The black area surrounded by colored area is the foreground information to be further processed.

### Identification of coarse background

To identify the approximate level of the ground noise for downstream fine thresholding, a general strategy is to define a highly representative “feature value” of ground noise, based on which a global threshold can be estimated to segment the “coarse background”. Guided by this strategy, we designed an algorithm that calculates a global threshold using the morphological features of the ground noise – namely, non-connected negative element scanning (NNES; Fig. 2b). In brief, NNES scans the total number of non-connected negative elements at different global thresholds, resulting in the identification of the representative value (i.e., the peak) with a maximum non-connected negative element count (Fig. 2b, (i)). The global threshold for the coarse background (Fig. 2b, (iii)) will be determined using a linear model trained by the representative value rendered by NNES and manual global thresholding (MGT), a global threshold determined by researchers experienced in manual cytoskeleton image analysis (Supplemental Fig. 1 and 2). NNES can maintain stability and accuracy over different samples that vary in the brightness distribution, because ground noise is the image component with the lowest value that is subject to a normal distribution and generally does not interfere with the actual fluorescence signal. Another accessible method is to directly use the peak-of-frequency brightness of the image as a representative value to train a model. However, this approach is less accurate because the interval near the theoretical peak is always turbulent and non-monotone, a limitation potentially due to the pollution of diffracted light (Supplemental Fig. 1; also see *Discussion*).

### Cytoskeleton segmentation by ILEE

The core strategy of ILEE is to accurately identify the edge of all cytoskeletal filament components and apply an implicit Laplacian smoothing^22^ on the selected edge. This leads to the generation of a threshold image where high gradient areas (i.e., the edge of cytoskeleton filaments) smoothly transit to each other (Fig. 3a). As illustrated in Supplemental Fig. 3a, the general local threshold trend changes as a function of the baseline values of the cytoskeleton edges. This is because ILEE selectively filters out high-frequency noise while preserving salient geometric features of individual actin filaments by leveraging the spectral characteristics of Laplacian operators (Supplemental Fig. 3b,c). Thus, ILEE can filter the background and noise regardless of the general local brightness level, demonstrating that its function does not require an operating kernel which would otherwise restrict the performance when evaluating filaments of varying thickness. Moreover, the edge of the cytoskeletal component can be smoothed and elongated using a significant difference filter (SDF; Supplemental Fig. 4) and a Gaussian filter, the sum of which serves to enhance the continuity of the edge and contributes to the accuracy of edge detection (Fig. 1b and 3a). For this reason, we name our algorithm as indicated as “Implicit Laplacian of Enhanced Edge”

**Fig. 3:**
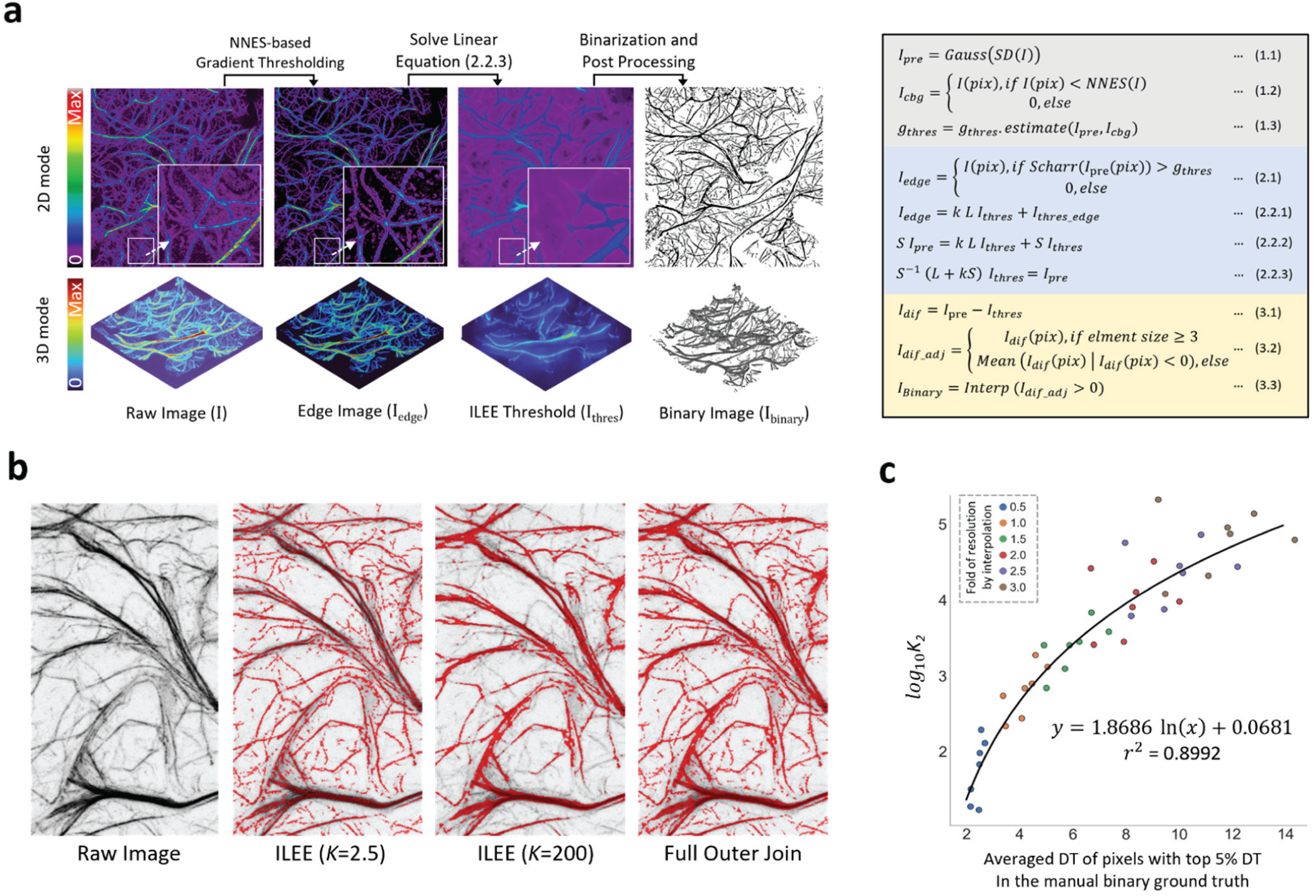
Cytoskeletal identification by ILEE. **a**, Visual demonstration and summarized mathematical process of ILEE. On the left, the visualized intermediate images of ILEE process are presented; on the right, an abbreviated mathematical process of ILEE is shown (grey, image pre-processing; blue, ILEE in a narrow sense; yellow, post-processing; also see *Method* for the detailed computational algorithm). **b**, The value of implicit Laplacian smoothing coefficient *K* influences ILEE performance. When *K* is small (e.g., 2.5), the rendering of faint and thin filaments is accurate, but the edge of thick filaments tends to be omitted. Conversely, when *K* is large (e.g., 200), rendering of thick filaments is accurate but thin and faint filaments are omitted. A solution is applied by creating a full outer join threshold image by a fixed *K*_*1*_ = 2.5 and an estimated universal *K*_*2*_ for an entire biological batch of samples. **c**, *K*_*2*_ estimation model. A regression model to compute any universal *K*_*2*_ for a given sample batch (see Supplemental Figure 8 for detail). To augment the training samples, 7 images with manually portrayed binary ground truth are interpolated by a new resolution of different folds to the original into 42 samples (shown as single dots) that cover the general range of actin thickness. A non-linear regression estimation model is trained using the highest 5% DT (distance transform) values in the ground truth (representing filament thickness) and an anticipated *K*_*2*_ with a specific satisfactory capability to detect thick filament (see Supplemental Figure 8 for detail). In the standard ILEE pipeline, the mean of top 5% DT of all input samples will be calculated by Niblack thresholding to obtain the universal *K*_*2*_ for ILEE.

For computation, the ILEE algorithm builds a linear system based on Laplacian operators to achieve local adaptive thresholding for edges of cytoskeletal components (see *Methods* for detail). In brief, we first used the information from the coarse background to estimate a global threshold of the gradient magnitude (*g*_*thres*_), and classified pixels/voxels above *g*_*thres*_ as boundary elements (*I*_*edge*_) (Fig. 3a, 2.1). Following the boundary identification, we constructed a *n* × *n* (*n* is the total number of pixel/voxel in the image) selection matrix *S*, a sparse diagonal matrix whose diagonal corresponds to the flattened pixel/voxel in order; the diagonal element was 1 if the pixel/voxel possessed a norm of the gradient above *g*_*thres*_. As the pixels/voxels belonging to the boundary of the cytoskeleton filament were marked, *I*_*edge*_ can be mathematically defined as shown in Fig. 3a, equations 2.2.1 and 2.2.2, where *L* is the Laplacian matrix and *K*, or implicit Laplacian smoothing coefficient, is a weight that adjusts the influence of the Laplacian operator. The local threshold image can therefore be rendered by solving the linear equation shown in Fig. 3b, 2.2.3.

For a given image input, the performance of ILEE depends on two parameters: *g*_*thres*_, which defines the edge, and *K*, which determines the weight of detail (i.e., high-frequency components) to be filtered. To calculate the *g*_*thres*_ for an input image, we used brightness values of the area identified as coarse background by NNES. Since the ground noise is approximately subject to a normal distribution, we hypothesized a deducible statistical relationship between the image gradient, defined by Scharr operator^23^, and the native brightness of pixels/voxels within the coarse background. Using a normal random array that simulates the noise with a 2D or 3D data structure, we demonstrate that the distribution of the background gradient magnitude is also normal-like, and both mean (*μ*_*G*_) and standard deviation (*a*_*G*_) of the gradients are directly proportional to the standard deviation of their native pixel values (*a*_*cbg*_), and we calculate the proportionality coefficient (see Supplemental Fig. 5). For the 3D mode, since the voxel size on the x- and y-axis is different from that of the z-axis, the proportionality coefficient of *μ*_*G*_ and *σ*_*G*_ over *σ*_*cbg*_ will vary for different ratios of the x-y unit : z unit (see Supplemental Fig. 6). To solve this problem, we simulated artificial 3D noise images and trained a multi-interval regression model that accurately (R^2^ > 0.999) calculates the proportionality coefficient of *μ*_*G*_ and *σ*_*G*_ over *σ*_*cbg*_ for different x-y : z ratios of the voxel. Finally, using this approach and randomly selected actin image samples, we trained a model, *g*_*thres*_ = *μ*_*G*_ + *σ*_*cbg*_ * *σ*_*G*_, to determine the *g*_*thres*_ as ILEE input (Supplemental Fig. 7).

To determine the appropriate setting of *K*, we first tested how different *K* values influenced the result of the local threshold image (*I*_*thres*_ of Fig. 3a). As shown in Supplemental Figure 8a, at the optimal *g*_*thres*_, a low value of *K* generated an *I*_*thres*_ that was highly consistent with the selected edge; when *K* increases, the total threshold image shifted towards the average value of the selected edges with increasing loss of detail. As for the resultant binary image, a lower *K* enables the accurate recognition of thin and faint actin filament components, yet cannot cover the entire width of thick filaments. Conversely, a high *K* value covers thick actin filaments with improved accuracy, resulting in a binary image that is less noisy; however, thin and/or faint filaments tend to be omitted as pseudo-negative pixels (Fig. 3b, Supplemental Fig. 8a). To overcome this dilemma, we applied a strategy using a lower *K*_1_ and a higher *K*_2_ to compute two different binary images that focus on thin/faint components and thick components, respectively. Then, we generated a full outer-join image that contains all cytoskeleton components in these two binary images. This approach led to improved recognition of cytoskeleton with varying morphologies (see Fig. 3b).

As described above, *K*_*1*_ controls the performance of thin and faint filaments. Since the theoretical minimum thickness of a distinguishable cytoskeletal component is approximately equal to one pixel/voxel unit, *K*_1_ can be fixed to a constant to recognize the finest cytoskeletal components from a complex and heterogeneous set of input samples. Using this approach, we identified an empirical optimal *K*_1_ of 2.5. However, since different image samples have different distributions of cytoskeleton thickness, *K*_2_, which controls the performance over thick filaments, must be guided according to the maximum thickness among all samples in an experiment batch. To ensure that the algorithm described herein is fully unguided, our strategy was to estimate an appropriate *K*_2_ from an estimated maximum thickness using all samples from a single batch of experiments, including multiple biological replicates (if applicable). To do this, we used Niblack thresholding first to generate a coarse binary image (which is sufficiently accurate for the thickest portion of the filament); from this, we calculated the mean of the top 5% of the Euclidian distance transformation (DT) values of all positive pixels (see *Methods* for additional information). Next, the top 5% means of all individual images were averaged to estimate *K*_2_ via a trained model using a sample set with manual portraited binary ground truth (Fig. 3c and Supplemental Fig. 8b, c, d). Therefore, individual images from all groups in the same batch of an experiment can be processed by *K*_2_ as estimated using the model above. In this, the bias of human input was avoided. When processing 3D images, we additionally provided a parameter that allows using a single *K*, rather than *K*_1_ and *K*_2_, which balances the accuracy over thin/faint and thick/bright filaments while saving certain amount of time of computation.

### Computational analysis of cytoskeleton indices

Through the image processing pipeline, cytoskeletal indices were automatically calculated from the binary image generated by ILEE. As a substantial expansion from three previously defined cytoskeletal indices (e.g., *occupancy, skewness*, and *CV*)^17,20^, we totally introduced 12 indices (Fig. 1a); particularly, we focused on 9 of the 12 indices in this study, as they are normalized and ready for biological interpretation (see Supplementary Fig. 9 for visualized demonstration for each index). These indices fall within five classes – density, bundling, connectivity, branching, and directionality, which describe the quantifiable features and functions of cytoskeletal morphology and dynamics. It is worth noting that many of these indices require a certain level of image post-processing (e.g., oversampling) to further enhance the accuracy of the output. The detailed mathematical definition of each index is described in *Methods*.

We have defined a novel set of cytoskeletal indices to enable the measurement of certain cytoskeletal features. For the class “density”, we developed *linear density*, a feature that measures filament length per unit of 2D/3D space. For the class “bundling”, we developed two new highly robust indices, referred to as diameter by total DT (*diameter_TDT*) and diameter by skeleton DT (*diameter_SDT)*. Both of them quantify the physical thickness of filament bundles, in addition to the indirect indices *skewness* and *CV*, which estimate the relative bundling level based on the statistical distribution of fluorescence intensity. For the class “connectivity”, we defined *static severing activity*, which estimates the severing events within per length unit of the cytoskeleton. *static severing* activity assumes that a severing event generates two visible cytoskeletal filaments, which is distinguishable from filament depolymerization. This is an important consideration in terms of the biological behavior of the cytoskeleton, as it enables the decoupling of the impact of filament depolymerization and filament severing, key activities facilitated by the eukaryotic actin depolymerizing factor (ADF) and cofilin family of proteins^24^. For the class “branching”, our algorithm is based on Skan, a recently developed Python library for the graph-theoretical analysis of the cytoskeleton^25^. To further explore the relationship between filament morphology and the biological activity of branching, we specifically designed a novel index, *static branching activity*, which we define as the total number of additional branches emerging from any non-end-point node in per unit length of filament. In total, this index measures the abundance/frequency of cytoskeletal branching. Finally, our library supports estimating the level of directional cytoskeletal growth by the index *local anisotropy*, which measures how local filaments tend to be directional or chaotic. This approach is transplanted from an ImageJ plug-in FibrilTool^26^, but we expanded this algorithm to 3D space (Fig. 1c).

It is noteworthy that we named the two indices describing severing and branching above as “*static” severing activity* and “*static” branching activity*, because, by the most rigorous mathematical criteria, a dynamic status cannot be directly defined without introducing the time dimension. However, the introduction of time dimension (using a video sample) generally sacrifices spatial resolution due to the technical limit of microscopy devices. While it is available to use TIRF (total internal reflection fluorescence microscopy) or VAEM (variable-angle epifluorescence microscopy) to collect 2D videos^27^, such methods may be limited by the insufficient quantity of filaments in the field of vision and manual counting of the dynamic events, both of which may introduce bias compared to a larger 3D data structure. Alternatively, we introduce *static severing activity* and *static branching activity* as indirect indices measuring the intensity of severing and branching. Here, we assume that the occurrence of severing/branching events and the net regeneration speed of cytoskeleton has established (or is very near to) their dynamic equilibrium; at this point, the quantity of pieces/branches of the filament is positively correlated to the speed of severing and branching. In practice, most experimental schemes set an interval between the treatment (if any) and imaging, which is long enough to reach the dynamic equilibrium. Therefore, we expect these two novel indices can provide interpretable and reliable data to estimate the intensity of severing and branching, especially for confocal images.

### ILEE displays high accuracy and stability over actin image samples

To evaluate the performance of ILEE in terms of its accuracy and compatibility over diverse samples, we selected a dataset of actin images from Arabidopsis leaves with diverse morphology, and compared ILEE with numerous traditional global and local thresholding algorithms, including MGT. First, to evaluate the accuracy of each algorithm in terms of filament segregation, we manually generated the ground truth binary image from each of the samples using a digital drawing monitor (Fig. 4a, ground truth). Next, we used each ground truth binary image as a reference and compared the filament architecture obtained by ILEE, MGT, and 6 additional adaptive thresholding algorithms. These additional thresholding algorithms include Otsu^18^, Triangle^28^, Li^29^, Yan^30^, Niblack^19^, and Sauvola^31^ (Fig. 4). As an additional concern of rigor, we anticipated that pseudo-positive pixels may be obtained due to operator’s bias during the generation of each of the ground truth images (even when the operator is experienced in the actin imaging field). Therefore, we further analyzed each non-connected component of pseudo-positive pixel by its shape and connectivity to the matched elements and identified the actin-like pseudo-positive pixels as possible real actin components.

**Fig. 4:**
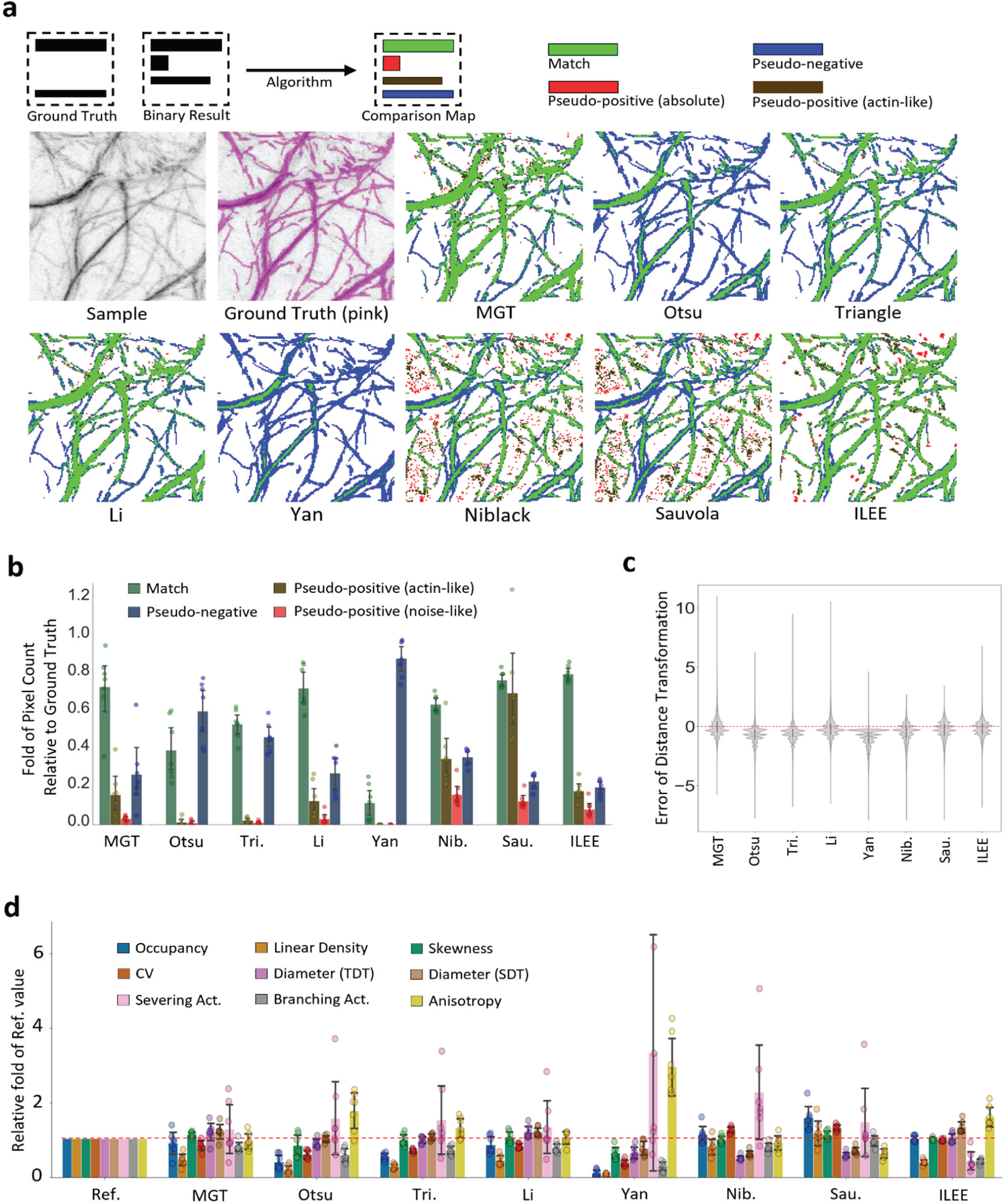
ILEE shows superior accuracy and stability over classic thresholding approaches. The manually portrayed binary ground truth of 7 visually diverse samples were compared with binary images rendered by ILEE, MGT, four global thresholding algorithms (Otsu, Triangle, Li, Yan), and two local thresholding algorithms (Niblack and Sauvola). **a**, Visualized comparison of ILEE versus other approaches. Pixels with different colors are defined as green: match of rendered binary image and ground truth; blue: pseudo-negative, the pixels omitted by the algorithm; red, noise-like pseudo-positive, the pixels that are rendered by the algorithm but not in the ground truth without a shape of filament; brown, actin-like pseudo-positive, the pseudo-positive pixels within a filament-like component, which cannot be judged by high confidence. ILEE has the most accurate cytoskeleton segmentation. **b**, Quantitative comparison of pixel rendering demonstrated that ILEE has superior accuracy and stability across diverse samples. ILEE has the highest match rate with low and stable error rate; MGT and Li also have acceptable performance. **c**, Comparison of distribution of distance transformation error. Single pixel errors of all 7 samples were merged and summarized as a violin plot. Red dashed line indicates no error, or results identical to the ground truth. ILEE has a symmetric and centralized distribution, indicating an accurate and unbiased filament segmentation. **d**, ILEE has accurate and stable computation of cytoskeletal indices. Nine biologically-interpretable cytoskeleton indices computed using binary images rendered by different algorithms were compared with the ground truth. The index values were normalized to the fold of ground truth. Red dashed line indicates 1-fold, or identical to the ground truth. Additional results are shown in Supplemental Figure 10-13.

As shown in Fig. 4a (visualized demonstration), 4b (quantitative analysis), and 4c (bias analysis), ILEE offers superior performance, with improved accuracy, reduced pseudo-positive/negative occurrence, and the lowest bias over local filament thickness compared to current approaches. It is noteworthy, however, that the adaptive global thresholding approaches (from Otsu to Yan) tend to be relatively accurate when judging the thick and bright bundles of the cytoskeleton. However, these approaches are unable to capture faint filaments, and as a result, generate a high pseudo-negative rate. Conversely, both adaptive local thresholding approaches, Niblack and Sauvola, generate numerous block-shaped pseudo-positive elements, and fail to capture the near-edge region of thick filaments. For MGT and Li method, although they showed satisfactory match rate and lower averaged pseudo-positive/negative rates, their performance is far less stable than ILEE (Fig. 4b).

As a next step, we evaluated the accuracy and stability of cytoskeletal indices using ILEE versus other commonly used imaging algorithms. To do this, we first computed the ground truth indices from the manually generated binary images; then, quantitative measurements were collected from all methods and normalized by the relative fold to the result generated from the corresponding ground truth image. As shown in Fig. 4d, ILEE showed the best overall accuracy and cross-sample stability compared to all other quantitative approaches, the highest accuracy for *occupancy, skewness*, CV, and *diameter_TDT*. However, we did observe that in terms of the morphology-sensitive indices (i.e., *linear density, severing activity*, and *branching activity*), the ILEE algorithm did not fully conform with data collected from the ground truth binary images. Upon further inspection, we determined that this is because the manually portrayed ground truth images and ILEE results showed different tendencies in judging the pixels in the narrow areas between two bright filaments (see *Discussion*). While other approaches displayed obvious, somewhat predictable, inaccuracies, the MGT and Li methods still generated satisfactory results, which echoes their performance in actin segmentation. However, the performance of these two algorithms among diverse and complex biological samples was not as stable as ILEE.

In order to further evaluate the stability and robustness of ILEE performance, we continued to analyze the variance coefficient of all groups (Supplemental Fig. 10), uncovering that ILEE is the only approach that simultaneously maintained high accuracy and stability. Next, we tested the robustness of ILEE and other approaches against noise signal disturbance by adding different levels of Gaussian noise to the image dataset (Supplemental Figs. 11-13). Using this approach, we observed that ILEE is still the best performing algorithm, maintaining stable and accurate image binarization and cytoskeleton indices against increasing noise. Taken together, these results demonstrate that ILEE has superior accuracy, stability, and robustness over MGT and other classic automated image thresholding approaches in terms of both cytoskeleton segmentation and index computation.

### 3D ILEE preserves more information and gains higher fidelity over 2D ILEE

While the 2D ILEE generally outcompetes MGT and other commonly used automated algorithms (Fig. 4), the comparison of cytoskeleton segmentation approaches using the manually portraited ground truth is currently limited to 2D images, because it is extremely difficult and error-prone to select the “3D ground truth” voxel by voxel. However, as the 3D mode theoretically preserves information from the z-axis and renders higher accuracy, it is necessary to circumvent the challenge and verify these merits. Therefore, we turned to a different strategy using synthetic artificial 3D actin images with certain ground truth to investigate the performance of 3D versus 2D mode (Fig. 5a).

**Fig. 5:**
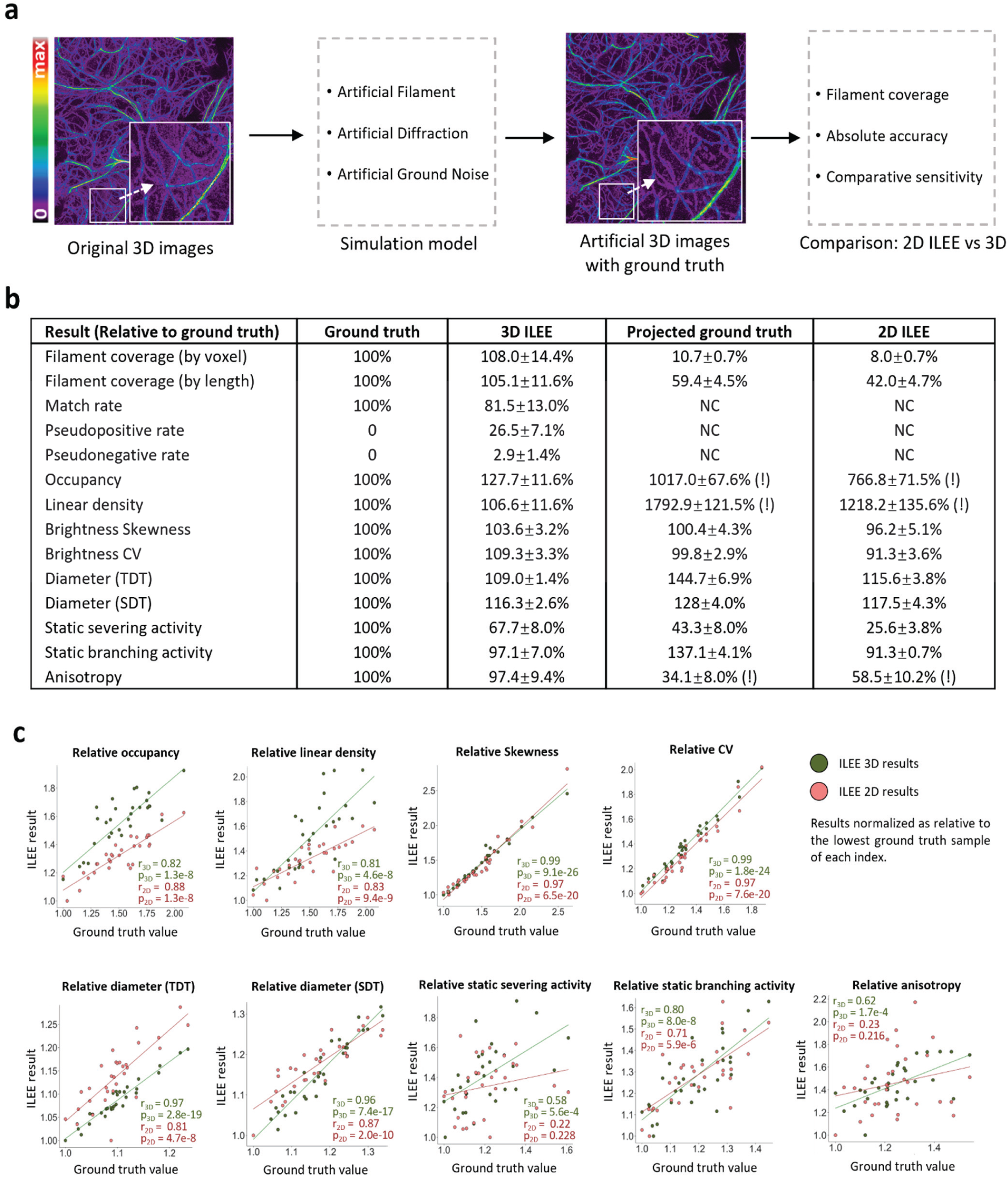
The 3D mode of ILEE renders cytoskeleton indices at much higher accuracy than the 2D mode, due to advanced data structure. We utilized image samples (N = 31, 25*800*800, Fig.2) to train an actin image simulation model that generates 31 artificial 3D actin images with ground truth. By comparing the performance of 3D and 2D ILEE over the artificial images, it was demonstrated that the 3D mode displays high fidelity to the ground truth while the 2D mode suffered from loss of information and systemic bias. **a**, The general experimental scheme. The raw 3D images are analyzed to train an actin image simulation model which uses statistical approaches to mimic the genuine actin filament, diffraction noise, and ground noise. Then 31 artificial actin images are regenerated with specific ground truth in terms of both actin segmentation and cytoskeleton indices. The 3D and 2D ILEE were applied to the artificial images, whose results were then compared to their corresponding ground truth. **b**, a table comparing the absolute accuracy of ILEE by the 3D and 2D mode. It was demonstrated that 2D mode losses ∼90% of pixels(voxels) and ∼50% of the total length of actin samples, and is less accurate for most of the indices. Exclamation mark (!): contrast difference due to different definitions of dimensional space in their units; not rigorously comparable. **c**, the linear correlation levels between ILEE results and the ground truth for all indices, by 2D and 3D mode. The absolute values of each index of the ground truth, 3D ILEE, and 2D ILEE are normalized by making their relative folds to the minimum data point of each group. A higher correlation coefficient r indicates a better capability to differentiate indices at different levels relatively, regardless of their absolute fidelity. The 3D mode has a generally better capability of differentiating high and low values.

First, we constructed an actin image simulation model, which mimics three different fractions of image brightness (real actin, ground noise, and diffraction noise; see Fig. 2a) using three independent statistical solutions (Supplemental Fig. 19; also see *Methods*). In brief, we utilized a training dataset of 31 diverse 3D samples to generalize the principles that describe the position-based brightness distributions of the voxels for the three fractions independently. Next, we adopted only the skeleton image generated by 3D ILEE from the training samples as the topological frame and refilled the whole image with new brightness values via the actin image simulation model. The advantage of this, rather than generating it *de novo*, was to ensure that the topological structure of the artificial filaments maximumly mimic the shape of genuine actin. Accordingly, we generated 31 3D artificial images and processed them using the ILEE 2D and 3D modes.

Our primary question was how 3D ILEE structurally improves the information loss through the 2D pipeline, and moreover, yields improved accuracy of the cytoskeletal indices. As shown in Fig. 5b, we observed that the 2D pipeline resulted in considerable information loss with reduced accuracy. Indeed, while the 3D segmentation almost accurately covers the filament voxels, the 2D pipeline only captured 8% of the data points as pixels and 42% of the total length of filaments, suggesting potentially biased sampling. Consequently, 3D ILEE generally gains index values closer to the ground truth than 2D ILEE, which indicates the 3D mode indeed possesses a better absolute accuracy/fidelity.

In some scenarios, studies are interested in the relative difference between experimental groups for biological interpretations, rather than the absolute accuracy of quantifiable features. Therefore, we also measured the “comparative sensitivity” of 3D and 2D ILEE, which reflects their capability to determine relatively high and low index values. To learn this, the index values of the ground truth images, as well as those computed through 3D and 2D modes, were first normalized to the fold to the minimum value of the corresponding group, followed by linear correlation analysis between the ground truth and ILEE outputs of each sample. As shown in Fig. 5c, the 3D mode has a higher correlation for seven of the nine indices, among which *diameter (TDT), static severing activity*, and *anisotropy* got a considerable improvement. Interestingly, we observed that the 2D mode performed slightly better for indices of the density class (e.g., *occupancy* and *linear density*). We posit that this is because z-axis projection may increase the contrast between samples with low and high filament density. In conclusion, our data suggest that the 3D mode of ILEE indeed eliminates the issues related to image projection and provides reliable results with higher accuracy and comparative sensitivity.

### ILEE leads to the discovery of new features of actin dynamics in response to bacteria

The primary impetus for creating the ILEE algorithm was to develop a method to define cytoskeleton organization from complex samples, including those during key transitions in cellular status. For example, previous research has demonstrated that the activation of immune signaling is associated with specific changes in cytoskeletal organization^27,32–34^. Complementary to these studies, other research identified the temporal and spatial induction of changes in the cytoskeletal organization as a function of the pathogen (e.g., *Pseudomonas syringae*) infection and disease development^35–37^. The sum of these studies, which broadly applied MGT-based quantitative analysis of cytoskeleton architecture, concluded that virulent bacterial infection triggers elevated density (by *occupancy*) yet induced no changes in filament bundling (by *skewness*) in the early stages of infection. Since one of our major motivations herein was to develop an ILEE-based toolkit – supported by novel cytoskeletal indices – to investigate the process of pathogen infection and immune signaling activation, we collected raw data from a previous study^34^ describing a bacterial infection experiment using Arabidopsis expressing an actin fluorescence marker (i.e., GFP-fABD2), followed by confocal imaging and data analysis by ILEE as well as MGT conducted by three independent operators with rich experience in actin quantificational analysis (Fig. 6). Additionally, because researchers sometimes apply a universal global threshold to all images from a batch of biological experiments to avoid tremendous labor consumption, we included this approach and aimed to observe its performance as well. In this experiment, the only categorical variant is whether sampled plants are treated with bacteria (EV) or not (mock). In total, nine indices that cover features of density, bundling, severing, branching, and directional order are measured and compared.

**Fig. 6:**
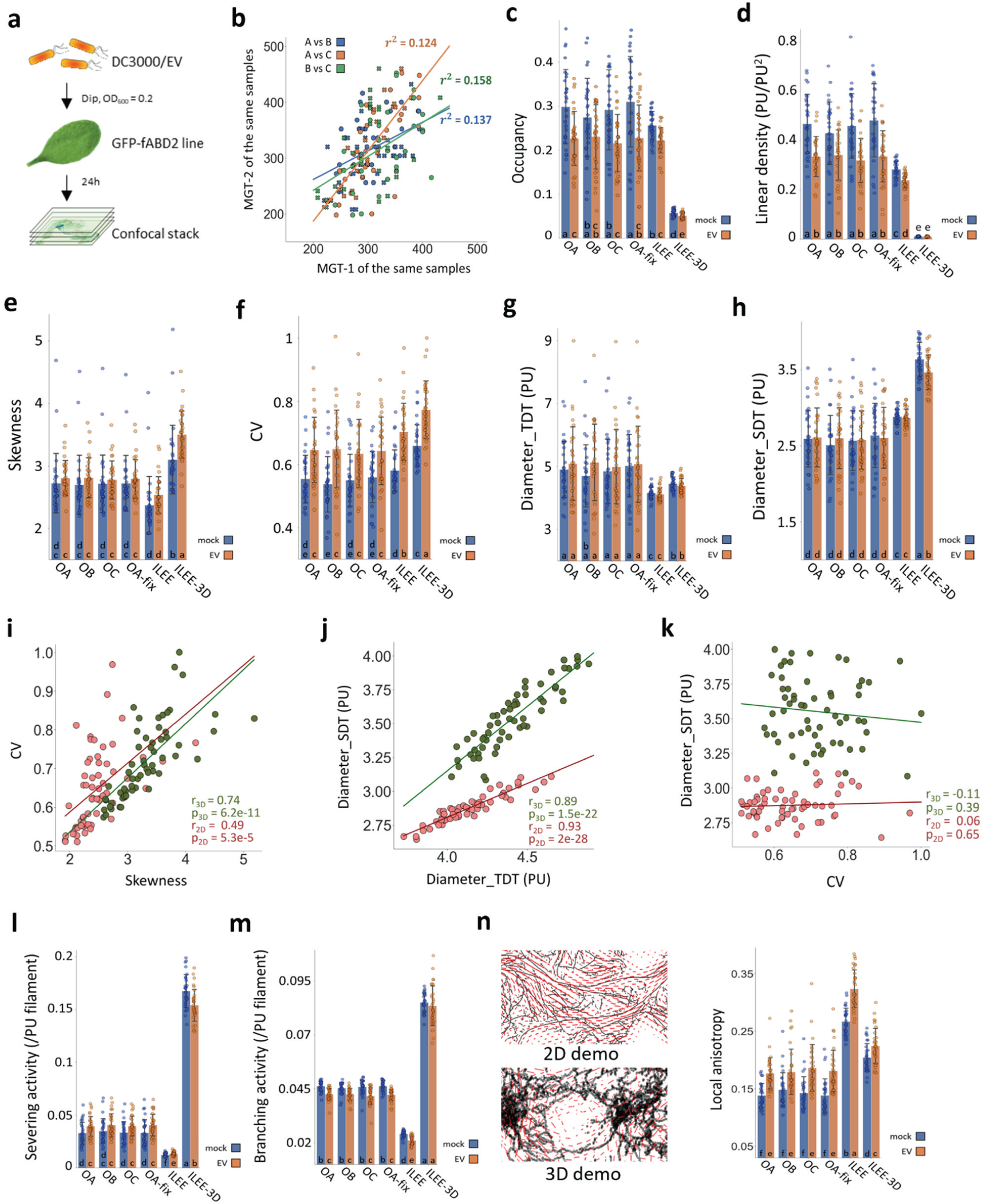
ILEE library enables the discovery of actin dynamic features of bacteria infected leaf tissue. Leaves of the Arabidopsis actin marker line Col-0/GFP-fABD2 were inoculated with mock or a virulent bacterial pathogen, *Pseudomonas syringae* pv. tomato DC3000 with empty vector (EV, identical to wild type); images (*n* = 28 for mock, *n* = 31 for EV) of epidermal cells were captured by laser scanning confocal microscopy at 24 hours post-inoculation (hpi). MGT and ILEE were applied to generate binary images and all indices were computed using ILEE_CSK. Double-blinded samples were provided to 3 operators (experienced researchers) OA, OB, and OC for comparison. OA additionally provides data using a universal threshold (OA-fix). **a**, Experimental schematic diagram. **b**, Corelative comparison of MGTs of individual samples determined by different operators. A very low correlation between each pair of operators indicates MGT has an increased risk of bias and is potentially inaccurate. **c, d, e, f, g, h, l, m, n**, Output indices *occupancy, linear density, skewness, CV, diameter_TDT, diameter_SDT, static severing activity, static branching activity*, and *local anisotropy*, respectively. Blue, mock; orange, EV. Multiple comparisons are conducted using a t-test without family-wise error correction (because image samples themselves are ground truth for all methods). Groups without overlapping letters have p_value lower than 0.05. For **n**, a visual illustration of the concept of local anisotropy is attached, where each red line segment points to the “averaged” direction of actin growth in the local area and the length shows the intensity of consistency of the direction. i, **j, k**, Comparisons of different indices in bundling class. *Skewness* and *CV* have medium-weak correlation; *diameter_TDT* and *diameter_SDT* have strong correlation; *diameter_SDT*, as a representative of direct indicator, and *CV*, as a representative of indirect indicator, have literally no correlation.

Our first question is whether the operator’s bias generated by MGT will influence the result and conclusion generalized from the experiment. We thereby analyzed the correlation of MGT results set by the three operators and found only a weak correlation between different operators (Fig. 6b), which indicates MGT indeed introduces bias and potentially impacts quantitative results. Interestingly, while minor statistical discrepancies between MGTs by different operators are found in some indices (i.e., skewness and severing activity), most of the MGT results (both adaptive or fixed) shows the same trend as 2D ILEE, yet with far higher standard deviation, or lower stability (Supplemental Fig. 14a) over a certain biological treatment. This indicates that the historical data based on MGT should be majorly trustworthy despite the biased single data points, but an accurate conclusion must be based on a high sampling number that hedges the deviation of individual data points. Since ILEE gains less systemic error over biological repeats, we also tested whether ILEE renders higher statistical power to identify potential significant differences. As suggested by Supplemental Fig. 14b, we that ILEE indeed has the lowest p-values among t-tests conducted for indices with a trend of difference. We believe these advantages demonstrate ILEE as a better choice for actin quantification.

Next, we attempted to understand whether different indices of the same class, particularly density and bundling, can reflect the quantitative level of the class in accordance, or instead show inconsistency. For density, we correlated the *occupancy* and *linear density* values of all methods over actin images of both mock and EV groups, and found that *occupancy* and *linear density* measurements are in high conformity, with a Pearson coefficient at 0.98 (Supplemental Fig. 15). For bundling indices, we were interested in their level of conformity because direct indices (based on binary shape) and indirect indices (based on brightness frequency distribution) are entirely different strategies to measure bundling. Using the same approach of correlation analysis, we found that *diameter_TDT* and *diameter_SDT* display strong positive correlation, while *skewness* and *CV* have merely medium-low correlation, which echoes the previous report demonstrating *skewness* and *CV* have different performance on the bundling evaluation^20^.

Unexpectedly, we also found that *CV* (as a representative of indirect indices) and *diameter-SDT* (as a representative of direct indices) have a striking zero correlation. This is perplexing, as it raises the question on whether *skewness* or *CV* should be regarded as an accurate measurement of bundling (see *Discussion*). This discrepancy is also observed by 3D ILEE, whose *CV* and *diameter-SDT* over mock versus EV revealed the converse results at significant differences. In general, we believe the biological conclusion that *Pst* DC3000 treatment renders increased actin bundling should be reconsidered with further inspection.

Lastly, we asked if additional features of plant actin cytoskeletal dynamics in response to virulent bacterial infection can be identified by the newly introduced indices and enhanced performance of ILEE. As shown in Fig. 6, we observed significantly regulated *static severing activity, local anisotropy*, and static *branching activity* triggered by the bacteria. At a minimum, these discoveries potentially lead to new biological interpretations, and as a result, may contribute to the identification of additional immune-regulated processes as a function of actin dynamics. However, while most 2D approaches were consistent and in agreement with the other indices, the *static severing activity* estimated by 3D ILEE indicates a significant but opposite conclusion. As suggested by the ground truth-based comparison on accuracy between 3D and 2D ILEE (Fig. 5), we believed this discrepancy was due to the information loss and misinterpretation by z-axis projection during the 2D pipeline. Hence, we have higher general confidence in the results and conclusion by the 3D mode.

### ILEE has broad compatibility with various sample types

Cytoskeleton imaging from live plant samples is arguably one of the most challenging types of images to evaluate due to the dynamic topology and uneven brightness of actin filaments. While we demonstrated that ILEE shows superior performance over plant actin samples, ILEE and the ILEE_CSK library are generally designed for non-specific confocal images of the cytoskeleton, therefore applicable to other types of samples. To investigate the compatibility of ILEE to other types of image samples, we tested ILEE on both plant microtubules^38^ and animal cell actin images (Supplemental Fig. 16). Importantly, we found that ILEE and ILEE_CSK library can process both plant and animal images with satisfying performance. This is encouraging, as ILEE can substitute or improve Hough transform, a straight-line detection algorithm commonly used for animal cytoskeleton (generally straight and thick), but with some limitation in neglecting and miscalculating curvy cytoskeleton fractions^14,15^. With the advancement of ILEE, Hough transform-based analysis may not be essential, and the potential cytoskeleton indices that rigorously require Hough transform can still utilize ILEE as a provider of binary image input for more accurate results.

## Discussion

We describe the creation of ILEE, an accurate and robust filament segregation algorithm for the unguided quantitative analysis of the organization of the eukaryotic cytoskeleton. As presented, this approach supports the *in vivo* analysis of both 2D and native 3D data structures, enabling an unbiased evaluation of cytoskeletal organization and dynamics. In addition to the development of key methods, we also generated a publicly available Python library that supports the automated computation of 12 filament indices of 5 classes of morphological features of the cytoskeleton. As described above, our data indicate that ILEE shows superior accuracy, robustness, and stability over existing cytoskeleton image analysis algorithms, including the widely adapted MGT approaches^17,39^. As a result of these newly developed approaches, we have developed an open-access library to conduct ILEE based cytoskeleton analysis, which eliminates limitations imposed by the classic 2D MGT approaches, including the introduction of user bias and different types of information loss.

### Robustness of NNES-based prediction of global gradient thresholds

The gradient threshold (*g*_*thres*_) defines the selected edge of actin filaments for implicit Laplacian transformation, the appropriateness of which greatly determines the performance of ILEE. To calculate *g*_*thres*_ without imposing a user-biased input, our strategy utilized specific feature values collected from the NNES curve and human-guided MGT to train a prediction model for the rapid rendering of a coarse area of the image background. Through this approach, we were able to deduce the corresponding *g*_*thres*_ by the mathematical relationship between the statistical distribution of the native pixel values and Scharr gradient magnitude of the coarse background (Supplemental Figs. 5, 6, 7, and *Methods*). This step might first appear unnecessary, since, alternatively, the most straight-forward strategy was to directly train a prediction model using the image gradient histogram and the human-determined *g*_*thres*_. However, as the gradient operator (see *Methods*) for any given object pixel is influenced by the values of surrounding pixels, the calculated gradient on the edge of the background is highly impacted (overestimated) by the contrast of foreground (cytoskeleton) versus the background. In other words, the brightness frequency distribution of the background gradient will change at elevated cytoskeleton fluorescence levels, even though the background *per se* does not. The outcome of this is a significant decrease in the accuracy of gained *g*_*thres*_. For this reason, we assert that *g*_*thres*_ should be mathematically deduced from a pre-determined region of background, rather than directly predicted via human-trained models or calculated from the frequency distribution of gradients even using the pre-determined background.

### ILEE and visual inspection have different inclinations on the topology between two bright filaments

Our data demonstrate that ILEE generally shows dominant robustness and accuracy for most indices, compared to the manually portrayed ground truth binary image. However, there are three indices (*linear density, static severing activity*, and *static branching activity*) where ILEE renders stable yet dramatically lower output than those derived from the ground truth images (Fig. 4d). Since the result is very stable, we anticipate there may be certain systemic inclinations in either the human-portrayed ground truth or the ILEE algorithm. After inspecting the binary images generated by ILEE compared with the ground truth, we identified a potentially critical reason: ILEE is less likely to presume the ambiguous, dubious thin connections inside the narrow space between two bright filaments. As demonstrated by Supplemental Fig. 17, while the ground truth and ILEE binary image look very similar, their skeleton images – which represent their topological structure – show a discrepancy between two bundles of bright filaments. Considering their procedure of generation, we speculate this is because: (1), ILEE fully outer-joins the binary results by a lower *K*_1_ and a higher *K*_2_ (Fig. 3c), among which *K*_2_ sacrifices the sensitivity to local filaments at the high signal region to improve the sensitivity at the low signal region, including the edge of thick filaments; and (2), human eyes tend to “hallucinate” imaginary filaments that do not statistically exist. According to our knowledge, there is no overwhelming evidence suggesting either 2D ILEE or the human eye is more accurate, but 2D ILEE is indeed more stable and conservative. However, 3D ILEE may solve this paradox because many “adjacent” bright bundles are artifacts out of z-axis projection, which is distant enough in 3D space to offer ILEE with a satisfactory resolution to split.

### The 2D and 3D modes of ILEE may lead to different conclusions

For the analysis over bacteria treated leaf sample (Fig. 6), 2D ILEE generally agreed with MGT while offering higher robustness and stability. Interestingly, this is not always the case for 3D ILEE, since it led to different statistical conclusions from 2D approaches in *linear density, skewness, diameter (SDT), static severing activity*, and *static branching activity*. However, such difference is understandable, because the 3D mode indeed computes indices with higher accuracy and overcomes many limitations by the 2D data structure. As demonstrated by Fig. 5b, c, 3D ILEE provide more or less improved absolute fidelity and comparative sensitivity for *skewness, diameter (SDT), static severing activity*, and *static branching activity*; therefore, our current data support that 3D ILEE is more trustworthy for these indices. Similarly, the discrepancy of *linear density* over 2D approaches versus 3D ILEE in Fig. 6d also agrees with our observation that 2D ILEE has higher comparative sensitivity for indices reflecting density. Hence, in terms of any potential discrepancy between the 2D and 3D modes, we offer a general recommendation: for occupancy only, 2D ILEE has better capability to distinguish biological difference; for all other cytoskeleton features, 3D ILEE is more trustworthy and dependable. Although it is currently difficult to make a definite conclusion whether the current 2D or 3D mode is more accurate across all scenarios, it is predictable that 3D computation will gradually substitute the classic 2D data structure, with the mainstream personal computers gaining increasing computational power in the future.

### Future development and applications

While ILEE has already remedied many disadvantages of traditional methods such as MGT, we are still working to advance further the ILEE approaches presented herein. Our goal is to ultimately establish algorithms and the toolbox that provide cytoskeletal indices at perfect accuracy and effectively answer the specific demands in this field of study. As such, we offer the following as a list of potential upgrades and applications to be integrated into the library:

- Deep learning-based cytoskeleton segmentation algorithm with “foreign object” removal function. As presented herein, ILEE enables the generation of trustworthy binary images on large scale, which enables the construction of deep learning models to identify cytoskeleton components from confocal images with potentially better performance. The deep learning-based approach is also the key to solving the ultimate problem of all current cytoskeleton segmentation algorithms (including ILEE), which is the inability to detect and erase non-cytoskeleton objects with high fluorescence, such as the nucleus and cytoplasm. As one approach to circumvent this limitation, free fluorescence proteins can be used as a cell-permeable false signal. This will enable us to train the model to recognize and exclude the non-cytoskeleton-like foreign objects, to render ideally pure cytoskeletal datasets.
- Vectorized intermediate image. After generating the difference image (i.e., *I*_*dif*_, Fig. 3a) using ILEE, one computational limitation of our Python-based algorithm is the tradeoff between the demand for unlimited high-resolution imaging versus limited computational power. Accordingly, an ideal strategy is to transfer the pixel/voxel image to a 2D vector image or 3D polygon mesh for index calculation. This will further enhance the accuracy of ILEE at an acceptable requirement of computational power.
- Regions of interest and organelle segmentation. There is currently a high demand in research of plant cell biology to quantify cytoskeletal parameters in stomatal guard cells, as well as additional cellular and subcellular architectures. In future releases of ILEE, we will develop approaches to enable automated recognition and selection of regions of interest, such as stomata, for various demands by the community.
- Compatibility to *x-y-t* and *x-y-z-t* data, where *t* represents time. We are in the process of developing a 4D-compatible analysis of cytoskeletal dynamics that tracks filament organization over time. This approach will provide a temporal evaluation of supported indices with high accuracy and robustness.

## Methods

### Plant genotypes and growth

*Arabidopsis thaliana* Col-0 expressing the actin cytoskeletal marker GFP-fABD2^36^ was used in this study. Arabidopsis seeds were stratified for 2 d in the dark at 4°C then sown onto the soil. All plants were grown in a BioChambers model FLX-37 walk-in growth chamber (BioChambers, Manitoba, Canada) at 20°C under long-day conditions (16 h of light/8 h of dark) with 60% relative humidity and a light intensity of approximately 120 μmol photons m^-2^s^-1^.

### Bacteria growth and plant inoculation

*Pseudomonas syringae* pv. *tomato* DC3000 (*Pst* DC3000) strains were grown as previously described^36^. Antibiotics used in this study include kanamycin (GoldBio #K-120-5, 50 μg mL^-1^), and rifampicin (GoldBio #R-120-1, 100 μg mL^-1^). Bacterial treatments for actin dynamics analysis were conducted following previously described methods^36^. Briefly, 2-week-old Arabidopsis Col-0/GFP-fABD2 was dipped for 30s in Dip-inoculation solution (10 mM MgCl2 + 0.02% Silwet-77) with DC3000 with empty vector (EV) at a concentration OD_600_ = 0.2 (ca. 2 × 10^7^ colony forming units/mL). Confocal images were collected at 24 hours post-inoculation.

### Mouse cancer cells sample

Yale University Mouse Melanoma line YUMMER1.7D4 cells (EMD Millipore, #SCC243) were cultured in DMEM (ATCC # 30-2006) supplemented with 10% FBS (Gibco #10437-028), 1% Pen-strep (ThermoFisher, #15140122), and 1% NEAA (Gibco, #11140035). For staining actin stress fiber, approximately 10,000 cells were seeded onto a glass coverslip maintained in a six-well plate overnight. First, semiconfluent cells were fixed with 3.7% formaldehyde for 15 min at RT, washed three times with PBS, then blocked in PBS supplemented with 2% BSA and 0.1% Triton for one hour at RT. Next, cells were incubated with 100 nM rhodamine-phalloidin (Cytoskeleton Inc, #PHDR1) in blocking buffer for 30 minutes at RT in the dark, then washed with PBS three times each for 5 min with gentle shaking at RT. Stained cells on the coverslip were mounted in ProLong Glass Antifade Mountant with DAPI (ThermoFisher, #P36982). Slides were cured overnight at room temperature and were then imaged.

### Confocal microscopy

For plant leaf actin images, 2-week-old Col-0/GFP-fABD2 plants were used for data collection and analysis. Images of epidermal pavement cells and guard cells were collected using laser confocal scanning microscope (Olympus FV1000D) by obtaining z-series sections at 0.5 μm intervals. Optical setting: 65x/1.42 PlanApo N objective with a 488 nm excitation laser and 510-575 nm emission filter. Images were collected at a resolution of 800 × 800 × 25 (*x*-*y*-*z*) and a 12-bit dynamic range.

Voxel size was 0.132 μm at the *x*- and *y*-axis and 0.5 μm at the *z*-axis. For animal cell actin images, YUMMER1.7D4 stained by rhodamine-phalloidin were sampled by the same confocal system. Optical setting: 100x/1.40 UPLSAPO objective with a 559 nm excitation laser and 570-670 nm emission filter. Images were collected at a resolution of 800 × 800 × 10 (*x*-*y*-*z*) with voxel size of 0.039 μm at the *x*- and *y*-axis and 0.16 μm at the z-axis

### Manually portrayed ground truth binary image

Seven raw projected images of 400×400 pixel size with diverse visual appearance (e.g., actin density, shape, thickness, fluorescence brightness) were selected from our actin image database. Using a pen-drawing display (HUION KAMVAS 22 Plus, GS2202) and GIMP, we enhanced the brightness of low-value pixels to clarify the actin structure and carefully portrayed the actin component of the selected image sample. The portrayed layer was extracted and transferred to binary format for further evaluation.

### Double-blind MGT analysis

For the mock versus *P. syringae*-inoculated sample pool, we erased the sample name and randomized the order and distributed them to three independent scientists with rich experience in cytoskeleton analysis (referred to as OA, OB, and OC) to let them determine the global threshold value of each sample manually using the approach described previously^35^. Once completed, we restored the grouping of the samples for batch analysis. We use Python to mimic the MGT pipeline generally conducted by ImageJ after determining a specific thresholding value. OA also provided a universal threshold value (referred to as OA_fix) that applies to all samples as a commonly used fast MGT approach.

### Statistical analysis and data visualization

All data analysis was conducted in the Python 3.8 environment and was described in figure legends.

### Determination of *K*_*2*_ for sample batches

For both 2D and 3D modes of ILEE, each 12-bit single-channel 3D image *I*(*x, y, z*) in a batch of samples was transferred into 2D image *I*_*proj*_(*x, y*) by z-axis maximum projection, where each pixel *I*_*proj*_(*x, y*) = max{*I*(*x, y, z*)|*z* ∈ *N*^+^, *z* < *z*_*max*_} · *I*_*proj*_ was processed by a Niblack thresholding function^16^ (library API reference [1]) to render *I*_*nibthres*_(*x, y*), with parameter *k* = -0.36 and *window_size* = 2 *int*(25 *l*) + 1, where *l* is the mean of x and y resolution of *I*_*proj*_. A binary image defined as *I*_*binary*_(*x, y*) = {1, *if I*_*proj*_(*x, y*) > *I*_*nibthres*_(*x, y*); 0, *else*} was generated. The binary image was processed through Euclidean distance transformation (library API reference [2]), and the mean of the highest 5% values was used as the input of the *K*_*2*_ calculation function (see Fig. 3) that outputs individual recommended *K*_*2*_. Finally, the mean of all individual *K*_*2*_ will be output as the recommend *K*_*2*_ for the total batch.

### ILEE

An abbreviated workflow of ILEE is illustrated in Fig. 3b. Here, we describe the overall process in further detail. For 2D mode, input image structure is *I*(*pix*) = *I*_*proj*_(*x, y*) = max{*I*(*x, y, z*)|*z* ∈ *N*^+^, *z* < *z*_*max*_}· For 3D mode, *I*(*pix*) = *I*(*x, y, z*). First, *I* is treated by a significant difference filter (SDF; Supplemental Fig. 4) to render *I*_*SDF*_, where:

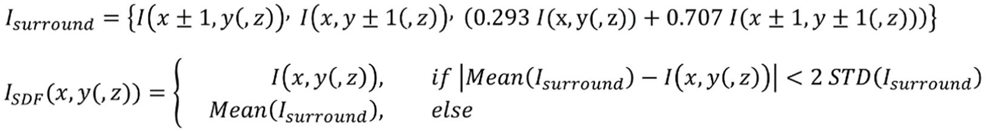

In other words, it substitutes a pixel by the mean of the 8 adjacent pixels if the absolute difference between it and that mean is higher than 2-fold of the standard deviation of the surrounding pixels. Then, *I*_*SDF*_ is input to a discrete gaussian filter with a 3×3(x3) weighting kernel at *σ* = 0.5, to render *I*_*pre*_ = *Gauss*(*I*_*SDF*_), which is the smoothed pre-processed image. Since confocal microscopy has different resolutions on x/y and z axis (hereby named as *U*_*xy*_ and *U*_*z*_), we adjusted the weighting kernel from *O*_*Gauss*_ to 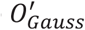 in 3D mode particularly by a scaling operator, as below:

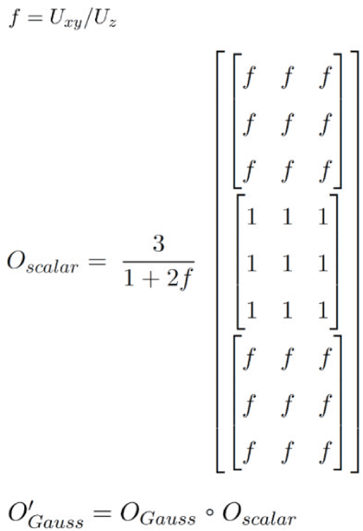

From *I*_*pre*_, the gradient magnitude image *G* is rendered through Scharr operator as:

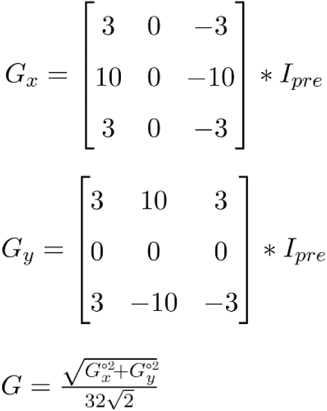

for 2D mode, or:

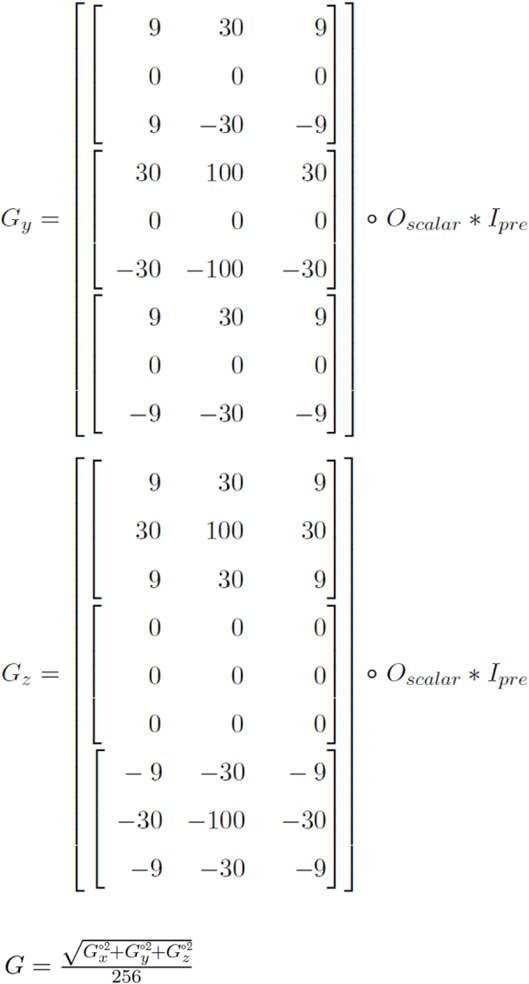

for 3D mode. Next, to calculate the gradient threshold (*g*_*thres*_) as an input for ILEE, a global threshold *t*_*cbg*_ to determine the coarse background (*I*_*cbg*_, as flattened image) is calculated by the NNES (non-connected negative element scanning) function that satisfies:

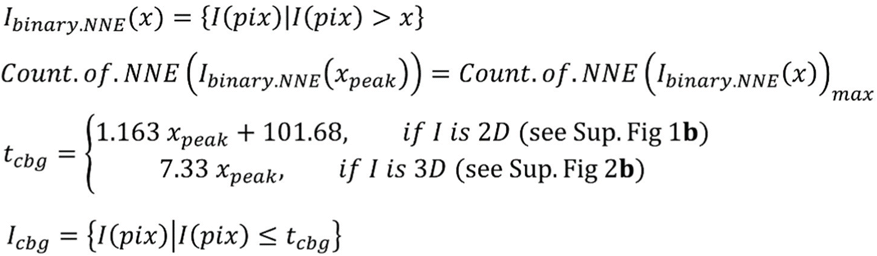

If *I* is processed using the 2D mode, as demonstrated previously (see Supplemental Figure 5), the statistical mean and STD of gradient magnitude of *I*_*cbg*_ (*μ*_*G*·*cbg*_ and *a*_*G*·*cbg*_, respectively) are univariate proportional functions of the STD of *I*_*cbg*_, *σ*_*cbg*_, not influenced by the mean of *I*_*cbg*_. Particularly, *μ*_*G*_ and *σ*_*G*_, of the coarse background area of the gradient image is:

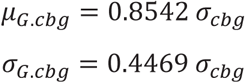

With these, we established a *g*_*thres*_ estimation model, where *g*_*thres*_ = *μ*_*G*·*cbg*_ + *k* (*a*_*cbg*_) *a*_*G*_· Using the optimized parameters described in Supplemental Figure 7, *g*_*thres*_ is deduced as:

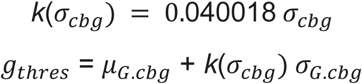

If *I* is processed by the 3D mode, the inconformity of *U*_*xy*_ and *U*_*z*_ will not only impact the weighting of Scharr operator, but also influencing the proportional coefficient of *σ*_*cbg*_ to *μ*_*G*_ and *σ*_*G*_. We simulated the accurate mathematical relationship between the proportional coefficient *k*_*s*_ and *U*_*z*_/*U*_*xy*_ (see Supplemental Figure 6), so *g*_*thres*_ for 3D can be calculated as:

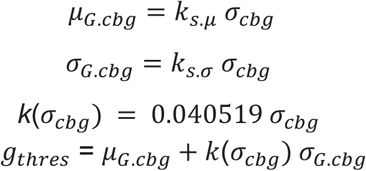

The total process above to computer *g*_*thres*_ is referred as function *g*_*thres*_· *estimate*(*I*_*pre*_, *I*_*cbg*_) in Fig. 3a (1.3).

The critical step to generate the threshold image of the input sample *I* is implicit Laplacian smoothing. This algorithm builds a linear system using the Laplacian operator to achieve local adaptive thresholding based on edges of cytoskeletal components. Leveraging the spectral characteristics of discrete Laplacian operators, we could filter out high-frequency components (aka, high fluorescence fractions of cytoskeleton) while preserving the low-frequency salient geometric features of background fluorescence. First, an edge image is defined as *I*_*edge*_(*pix*) = *I*_*pre*_(*pix*), *ifG*(*pix*) > *gthres*; 0, *else*, but we transform *I*_*edge*_ into a flattened image vector 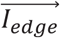, and hence it can be further be represented as:

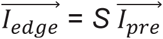

where *S* is a (sparse) selection matrix, which is a diagonal matrix with i-th diagonal entry being 1 if the i-th pixel has a gradient above *g*_*thres*_.

Next, according to the concept of Laplacian operator, we constructed the (sparse) Laplacian matrix *L* that satisfies:

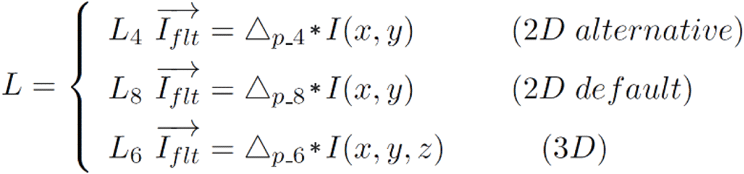

where the Laplacian operators in different modes are defined as:

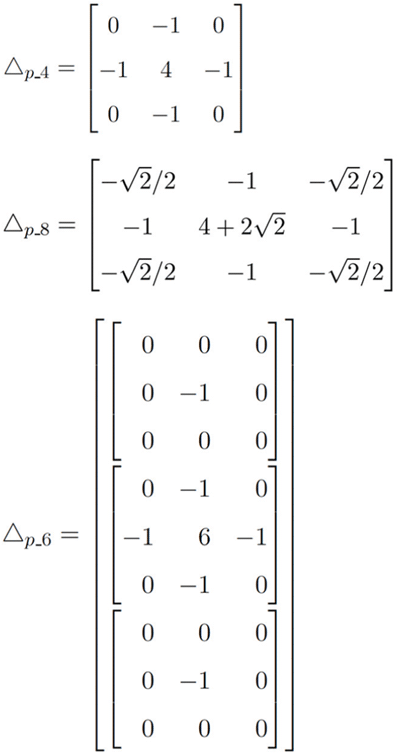

Therefore, we can establish an implicit Laplacian linear system that targets the edge:

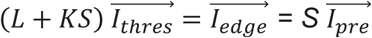

where 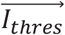 is the unknown to be solved by conjugate gradient method and restored to 2D/3D array *I*_*thres*_, and *K* is a weight that adjusts the influence Laplacian operator has on the result (see Fig. 3B).

Finally, a difference image *I*_*dif*_ is calculated as *I*_*dif*_ = *I*_*pre*_ − *I*_*thres*_. To further clean the noise, a temporary binary image *I*_*binary*·*temp*_(*pix*) = {1, *if I*_*dif*_(*pix*) > 0; 0, *else*} is generated and the sizes of all positive connected component (element) are counted. An adjusted difference image *I*_*dif_adj*_ is generated by lowering the values of potential noise into the mean of negative pixels:

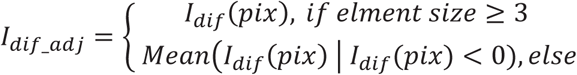

Using *I*_*dif_adj*_, the various versions of the binary image for computation of different indices are calculated as:

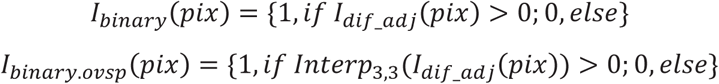

for the 2D mode, where *Interp*_3,3_ (*I*_*dif*_) is an oversampled image by bicubic interpolation whose resolution on both x-and y-axis are upscaled for 3 folds, or

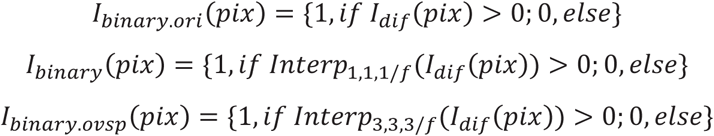

for the 3D mode, where *Interp*_1,1,1/*f*_(*I*_*dif*_(*pix*)) aims to restore the voxel to cubic shape by only interpolate z axis to 1/*f* folds of its original resolution and *Interp*_3,3,3/*f*_(*I*_*dif*_) additionally enhance 3 folds of resolution for all three axes.

### Computation of cytoskeletal indices

- ***Occupancy*:** the frequency of the positive pixels in the computed binary image, or ∑ *I*_*binary*_(*pix*)/*N* for 2D mode and ∑ *I*_*binary*·*ori*_(*pix*)/*N* for 3D mode, where *N* is the number of total pixels.
- ***Linear density*:** the length of the skeletonized filament per unit of 2D or 3D space. For 2D mode, *I*_*binary*·*ovsp*_ is skeletonized using Lee’s approach^18^ to render the skeletonized image *I*_*sk*_ · Then, *linear density* is calculated as:

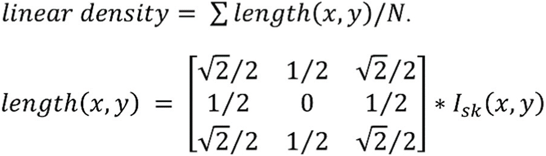

For the 3D mode, *I*_*sk*_ is rendered by *I*_*binary*_, and we use the sum of the Euclidean lengths of all (graph theory defined) branches obtained by Skan library^22^ as the total length of skeletonized filament and divide it by *N*. This is because sphere-cavities structures existent in the 3D skeletonized images are not applicable to the concept of length.
- ***Skewness*:** the (statistical) skewness of the fluorescence value of positive pixels, or mathematically:

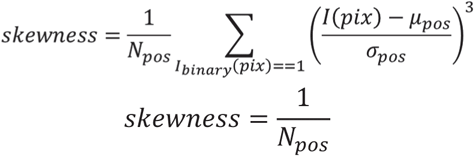

where N_pos_, μ_pos_, and σ_pos_ represents the count, mean, and standard deviation of positive pixels in the raw image *I*.
- ***CV*:** the (statistical) coefficient of variance of the fluorescence value of positive pixels, or mathematically:

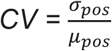

**Diameter_TDT**:average filament diameter estimated by Euclidian distance transformation of total binary image. The Euclidian distance transformation map *I*_dis_ is calculated as an image with the same shape of *I*_*binary*·*ovsp*_, but the value of each pixel is:

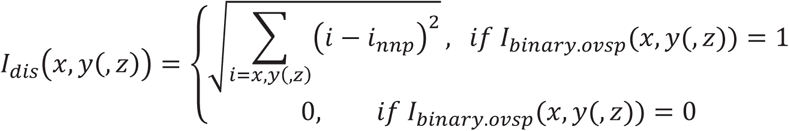

where *i*_*nnp*_ is the coordinates of the nearest negative pixel to (*X,Y,Z*). Therefore, the *Diameter_TDT* is mathematically defined as:

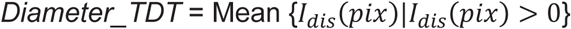
- ***Diameter_SDT*:** average filament diameter estimated by Euclidian distance transformation values of *I*_*binary*·*ovsp*_ *but samp*1*ed using on*1*y* positive pixels on *I*_*sk*_, or mathematically as:

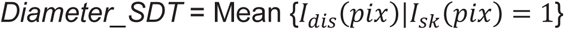
- ***Static severing activity*:** the count of connected components in the binary image per unit length of skeletonized filament. For images captured in 2D mode, it is *Count*· *of*· *NNE* (*I*_*binary*·*ovsp*_)/∑*length*(*x, y*). In 3D mode, both the count of non-connected elements and the length of skeletonized filament are called from the Skan library using *I*_*sk*_ computed from *I*_*binary*_ as the input.
- ***Static branching activity*:** the branching point count per unit length of skeletonized filament. *I*_*sk*_ is obtained from *I*_*binary*·*ovsp*_ for 2D mode or *I*_*binary*_ for 3D mode respectively and is next input into Skan library. The total number of type-3 and type-4 branches^22^ is collected and then divided by the length of the skeletonized filament.
- ***Local anisotropy*:** We performed a local averaging of filament alignment tensor, which is constructed as follows. First, we calculate the unit direction vector for each straight filament segment *g*_*i*_. Then, the covariance matrix for each segment is obtained from the following equation:

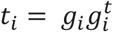

This rank-2 tensor is independent of the orientation of the line segment, and can thus be averaged over a region containing a collection of unoriented line segments. We weighed each filament tensor in a circular/spherical neighborhood by the length of every filament to produce a smoothed tensor field. The eigenvector corresponding to the largest eigenvalue indicated the primary orientation of filaments in this region. The difference between the maximum and the minimum eigenvalues is an indicator of the anisotropy in this region. If all the eigenvalues are the same, the indicator is 0, which implies an isotropic region. If the eigenvalues other than the maximum are all nearly 0, all the filaments in this region are parallel to each other. In this case, they are all aligned with the maximum eigenvector, the dominant filament direction of this region.

#### Simulation model of 3D artificial actin image

For each of the 31 training samples *I*(*x, y, z*), the binary image *I*_*binary*_ and skeleton image *I*_*sk*_ were previously obtained via ILEE, as described above. Three independent fractions reflecting artificial actin (*I*_*atf*·*a*_), artificial ground noise (*I*_*atf*·*gn*_), and artificial diffraction noise (*I*_*atf*·*d*_) are simulated independently.

To calculate *I*_*atf*·*a*_, the brightness of the real actin (*I*_*a*_) of all training sample is collected as:

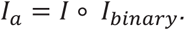

Here, we define the “reference voxel” of a non-zero voxel in *I*_*a*_ as its nearest voxel whose position has a positive value in *I*_*sk*_. In other words, for each non-zero voxel of *I*_*a*_(*x, y, z*), the distance from (*x, y, z*) to its nearest reference voxel (*P*_*R*_ = (*x*_*R*_, *y*_*R*_, *z*_*R*_)) on the skeleton image is defined as *D*_*R*_(*x, y, z*), where

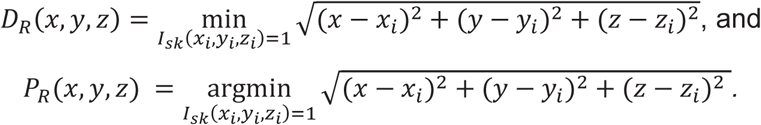

Meanwhile, the distance from the reference voxel to the nearest non-actin voxel, *D*_*R*2*N*_(*x, y, z*), is calculated as:

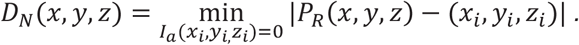

Next, the “relative eccentric distance”, *D*_*rlt*_(*x, y, z*), and the relative brightness comparing to the reference voxel, *I*_*rlt*_(*x, y, z*) are defined as:

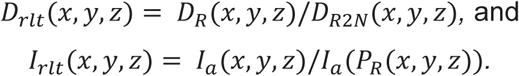

Note that *D*_*rlt*_ and *I*_*rlt*_ default to 0 whenever the denominator is 0. Because *D*_*rlt*_ is discrete, each possible value of *D*_*rlt*_ can be counted as n(*D*_*rlt*_). Using the *I*_*rlt*_ and *D*_*rlt*_ of all voxels with *D*_*rlt*_ ≤ 1 in the entire 31 images, a polynomial regression model between *D*_*rlt*_ and *I*_*rlt*_ is fitted using *log*_10_*n*(*D*_*rlt*_) as weights and adjusted into a piecewise polynomial function:

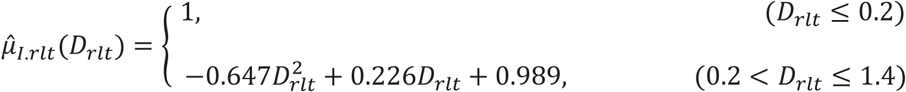

Similarly, the STD of the *I*_*rlt*_ is also fitted as below:

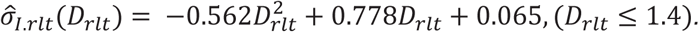

Finally, the artificial actin is generated as random values:

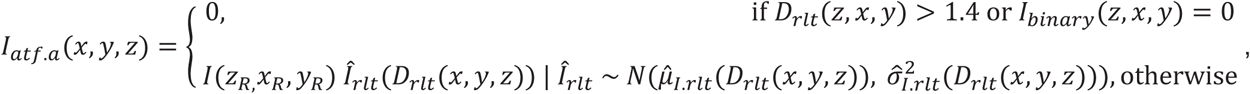

where *N* means normal distribution. The random values are clamped to [0,4095].

To calculate *I*_*atf*·*gn*_, two different strategies are utilized to generate two groups of voxels. For the voxels positions in the coarse background of *I*, we apply a fold of change that is subject to a normal distribution to each *I*(*x, y, z*) as the new ground noise, plus a Gaussian filter (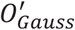, see ILEE section of *Method*) to neutralize the magnified gradient; for the rest beyond the coarse background, we gave random values that subject to a Burr-12 distribution (Library API reference [3]) fitted by voxels in the coarse background of *I*(*x, y, z*), that is:

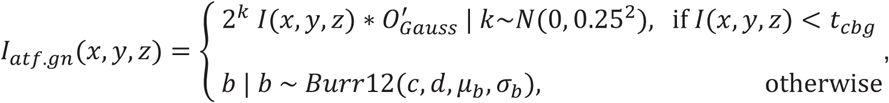

where *c, d, μ*_*b*_, and *σ*_*b*_ are the fitted parameters of the Burr-12 distribution using the training voxels {(*x, y, z*) | /(*x, y, z*) ≤ *t*_*cbg*_}. Similarly, the random values are clamped to [0,4095].

Third, for *I*_*atf_d*_, voxels of the positions in *I* that are neither coarse background nor real actin are used to fit an exponential distribution:

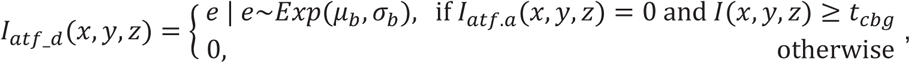

where *μ*_*e*_ and *σ*_*e*_ are the fitted parameters of the exponential distribution using the training voxels {(*x, y, z*) | *I*(*x, y, z*) ≥ *t*_*cbg*_}. Similarly, the random values are clamped to [0,4095].

Finally, these fractions of brightness are summed to generate the complete artificial image *I*_*atf*_, as:

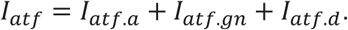

## Acknowledgements

We would like to thank Dr. Yi-Ju Lu (Michigan State University), and Dr. Silke Robatzek (Ludwig-Maximilians University - Munich) for generously providing raw images of actin and tubulin for experimental analysis and proof-of-concept analysis, respectively. We would like to thank Dr. Yi-Ju Lu and Dr. Masaki Shimono for providing individual MGT evaluations for our experiments. We would like to thank Rongzi Liu (University of Florida) for providing critical advice on statistical analysis and modeling. We would like to thank Dr. Richard Neubig (MSU) and Dr. Noel Day (Zoetis) for critical advice on manuscript preparation. We would like to thank Sarah Khoja for improving the reading experience of ILEE_CSK documentation website. Research in the laboratory of B.D. was supported by grants from the National Science Foundation (MCB-1953014) and the National Institutes of General Medical Sciences (1R01GM125743). Research in the laboratory of Y.T. is supported by grant from National Science Foundation (III-1900473).

## Author contributions

P.L., Z.Z., B.D., and Y.T. participated in the design and conception of ILEE pipeline and experiments. P.L. and Z.Z. developed the ILEE algorithm and ILEE_CSK library. P.L., Z.Z., and B.M.F. performed the experiments. P.L., Z.Z., B.D., and Y.T. analyzed and interpreted the data. P.L. and B.D. and wrote the manuscript. All authors actively edited and agreed on the manuscript before submission.

## Supplemental information

- S1. Supplemental Figures.pptx

