## Supplemental Figures for "Implicit Laplacian of Enhanced Edge: An Unguided Algorithm for Accurate and Automated Quantitative Analysis of Cytoskeletal Images"

[1] Department of Plant, Soil and Microbial Sciences, Michigan State University, East Lansing, Michigan 48824, USA. [2] Department of Plant Biology, Michigan State University, East Lansing, Michigan 48824, USA. [3] Department of Computer Science and Engineering, Michigan State University, East Lansing, MI 48824, USA. [4] Department of Pharmacology and Toxicology, Michigan State University, East Lansing, MI 48824, USA. [5] Molecular Genetics and Enzymology Department, National Research Centre, Dokki, Egypt.

*These two authors contributed equally to this work.

#### Supplemental Figures 1-18

#### Supplemental figure 1: NNES (Non-connected negative elements scanning) can locate coarse background better

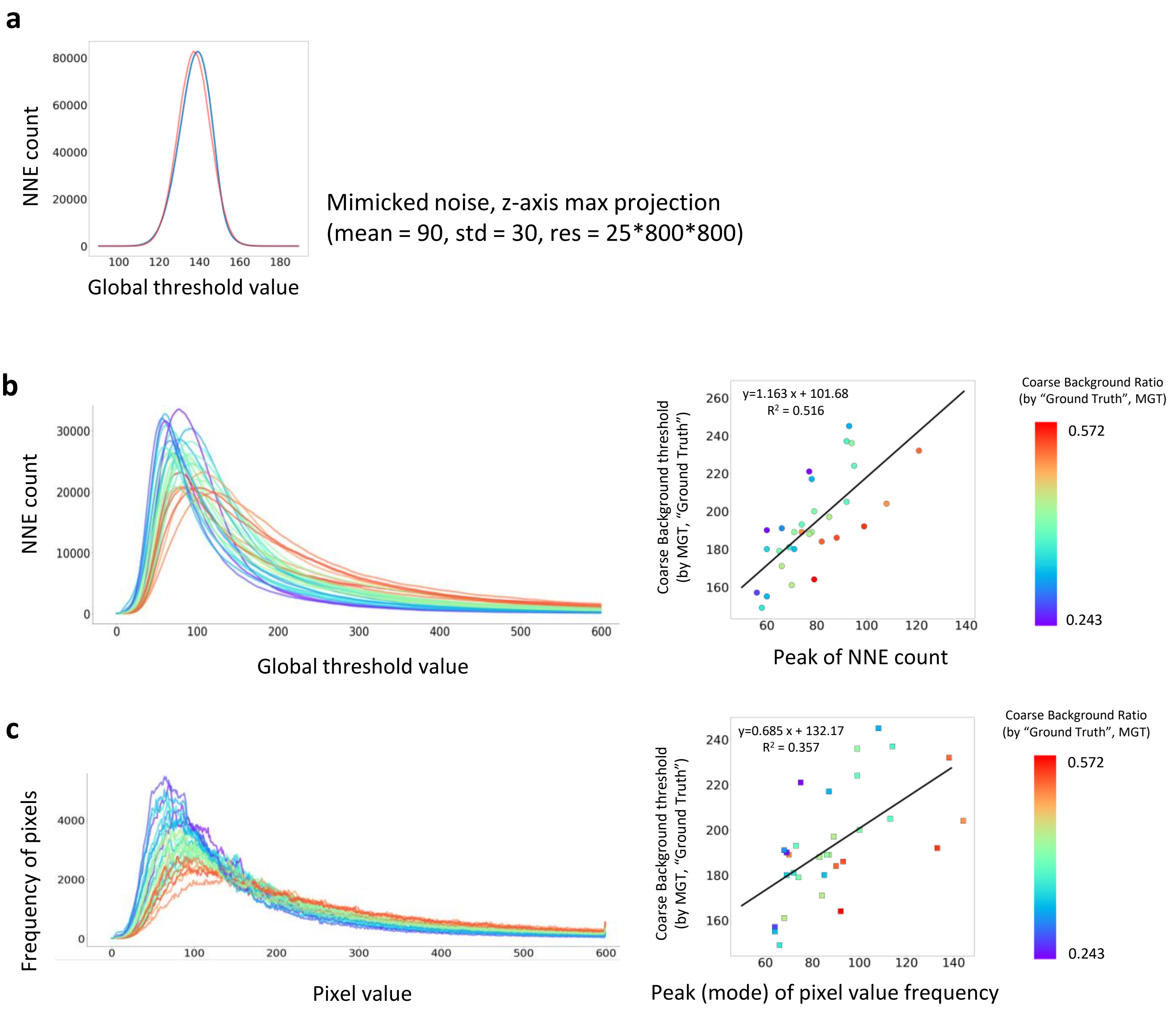


**Supplementary Figure 1. NNES (Non-connected negative elements scanning) identification of course background. a**, NNE count has a normal-like distribution. A confocal microscopy background-noise image was mimicked by random normal distribution (mean = 90, std = 30) into a 25*800*800 array. The maximum projection is conducted by choosing the maximum value of the third axis to make an 800*800 image (distribution shown as blue), and its mean and STD are calculated to make a true normal distribution (shown as red), both of which finely overlap. Therefore, the peak (representing the mean) is a good feature value. **b**, Performance of NNES-based adaptive global thresholding and its prediction. A random set of 31 actin image of 25*800*800 in our database are used to evaluate a coarse background threshold using MGT. Left, the NNES curves reflecting the relationship between global threshold and NNE count; right, correlation of ground truth coarse background evaluated by MGT vs peak of NNE count for individual samples. The color of each sample represents the proportion of coarse background within all pixels. **c**, Performance of brightness based adaptive global thresholding using histogram peak as a feature value. Left, histogram curve of the 31 training samples; right, correlation of the same ground truth coarse background vs peak of histogram (i.e., mathematical mode). Comparing to the histogram-based methods, NNES has a much smoother shape that enables the utilization of peak as a feature value. Also, NNES has a more robust correlation to coarse background determination, as by MGT.

**Supplemental figure 2: Performance of NNES adaptive global thresholding and its prediction model (3D)**

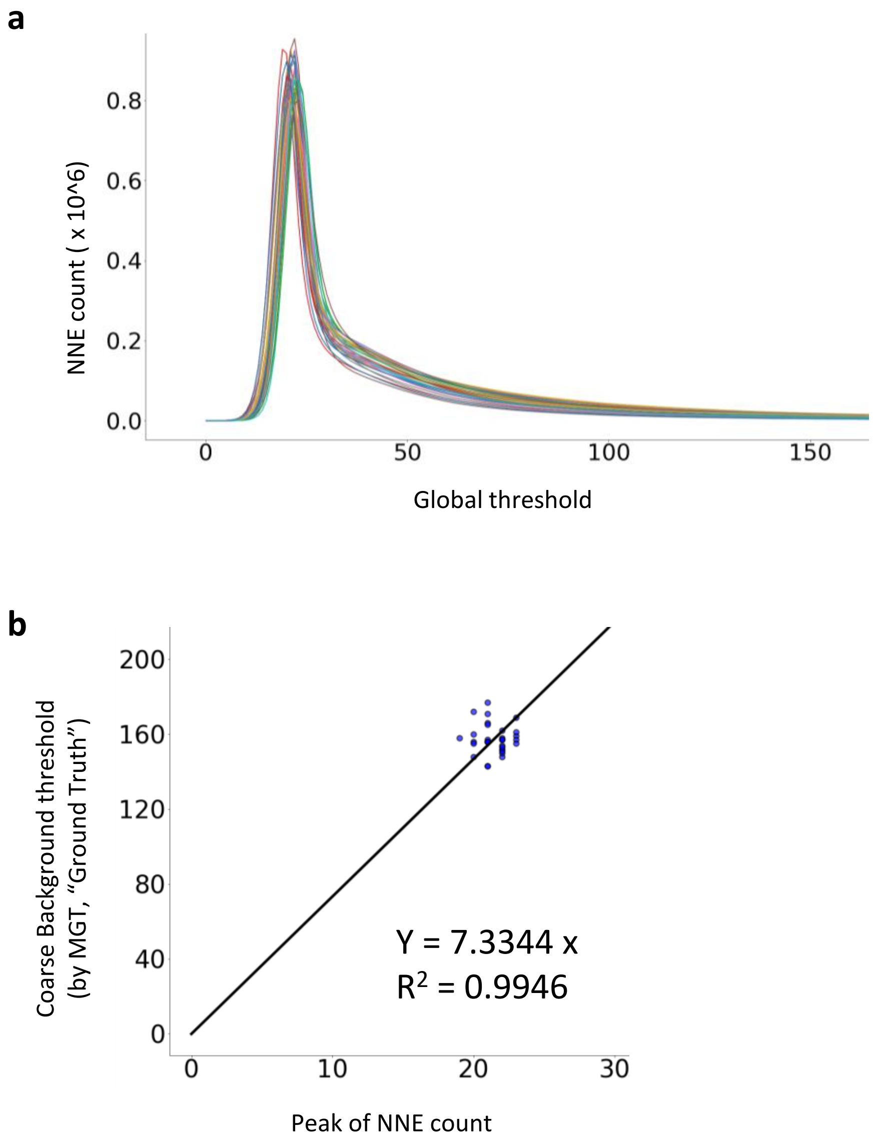


**Supplementary Figure 2. Performance of NNES adaptive global thresholding and its prediction model (3D). a,** The NNES curve of the same sample set as Supplement Figure 1. While they varies in actin features (density, bundling, etc.), they have very similar NNES shape. **b,** The correlation of coarse background (evaluated by an approach mimicking 2D MGT) vs the peak of NNE count for individual samples and corresponding coarse background prediction model. As suggested by NNES curve shape, they have very similar peak as well as coarse background, which represents the norm of sensor performance in 3D imaging (2D projection therefore is distorted by information loss). Therefore, we directly used a simple proportional function to establish the regression model.

#### Supplemental Figure 3: Visualized explanation of core ILEE algorithm

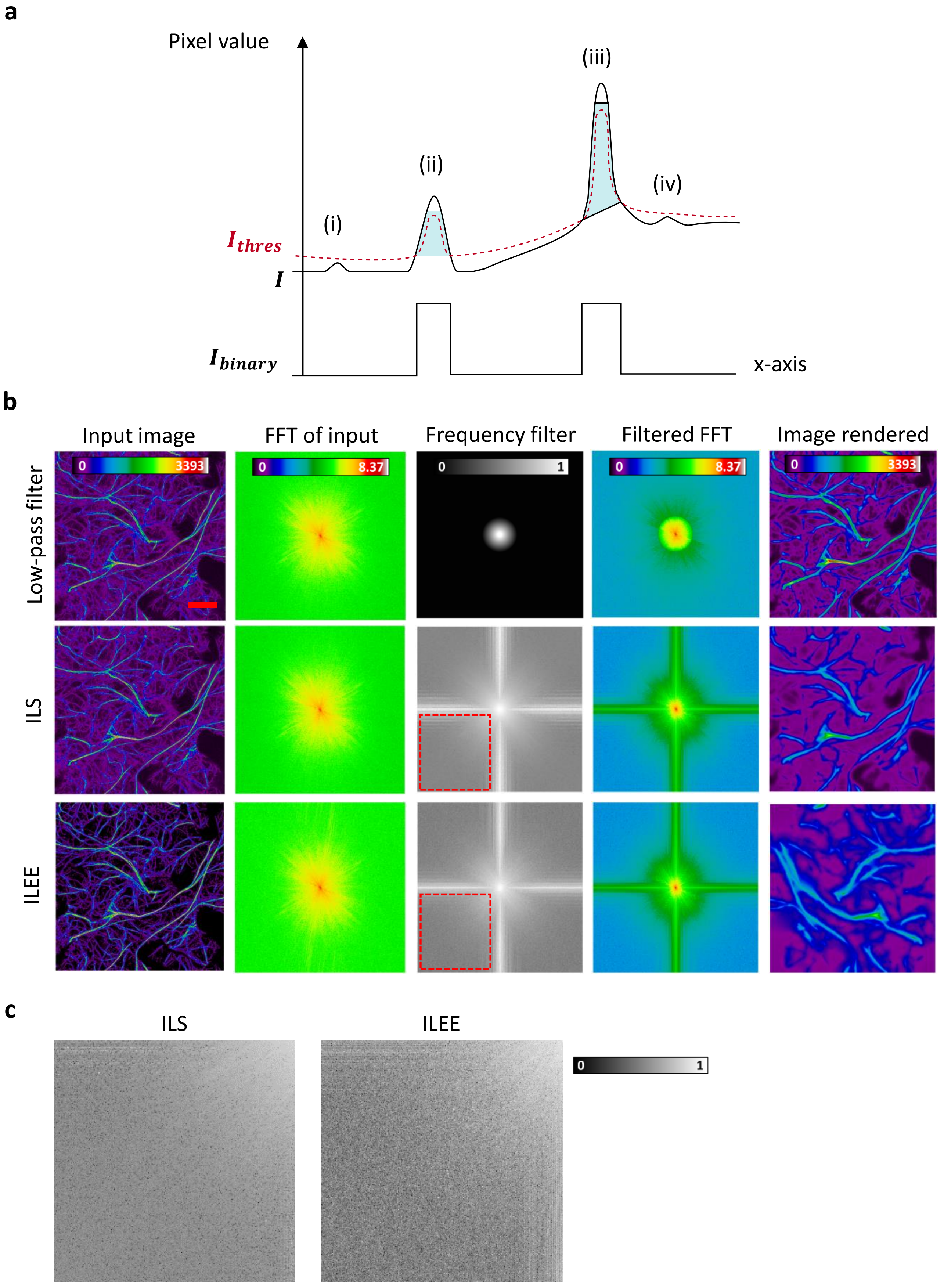

**Supplementary Figure 3: Visualized explanation of core ILEE algorithm.** The core strategy of ILEE thresholding is explained from the perspective of time domain threshold (marked cyan in a), which exclude(i.e., image itself) and frequency domain (i.e., after Fourier transformation, FFT). **a**, Schematic diagram describing how ILEE generates local threshold based on detected edges. For the purposed of simplicity, we use a "1D image" for demonstration. Suppose there is an example grayscale image ($\boldsymbol{I}$) with 4 peaks of pixel value (i, ii, iii, iv); the higher peak ii and iii are true cytoskeleton fluorescence but the lower peaks i and iv are random noise. ILEE first identify edges of cytoskeleton – area with a gradient magnitude higher than a computed threshold. Then these edge are used as "reference values" to generate a threshold image ($\boldsymbol{I_{thres}}$, in red color), that smoothly links all the reference area. Finally, the bona-fide cytoskeleton will be defined as area where $\boldsymbol{I}$ is higher than $\boldsymbol{I_{thres}}$. The areas not selected as edges will not be referred, which means true florescence with locally high values (ii, iii) are selected and background with locally low values (i, iv) are excluded, regardless of whether the local level is generally low (i, ii) or high (iii, iv). **b**, The comparison of effect of ILEE, classic low-pass filter, and implicit Laplacian smoothing (ILS) on image frequency domain. For low-pass filter, the input image (I) transformed into frequency domain pattern by FFT, and we artificially define a filter where 0-1 indicates the passing rate of each frequency fraction. We pass the frequency pattern through the filter to render the filtered FFT pattern, and restore the image by reverse FFT. For ILS and ILEE, the input image (I and Iedge respectively) are transformed to frequency domain pattern by FFT; on the other side, the result of ILS and ILEE are pre-computed and transformed to frequency domain pattern as the "filtered FFT". The (equivalent) frequency filter is deduced by subtracting filtered FFT from FFT of input and make the value relative to the FFT of input, which is comparable to the classic low-pass filter. ILS and ILEE has more fractions low frequency fractions on either x- or y-direction, which are potentially thick line cytoskeleton structures. Scale bar = 20 μm. **c**, Comparison of high-frequency filtering of ILS and ILEE. The red rectangle in subplot **b** is maximized to present the detail. ILEE has uneven but well-directed selection of high frequency fractions because ILEE particularly preserves the cytoskeleton edge and tends to neglect the coarse background.

#### Supplemental figure 4: Significant difference filter.

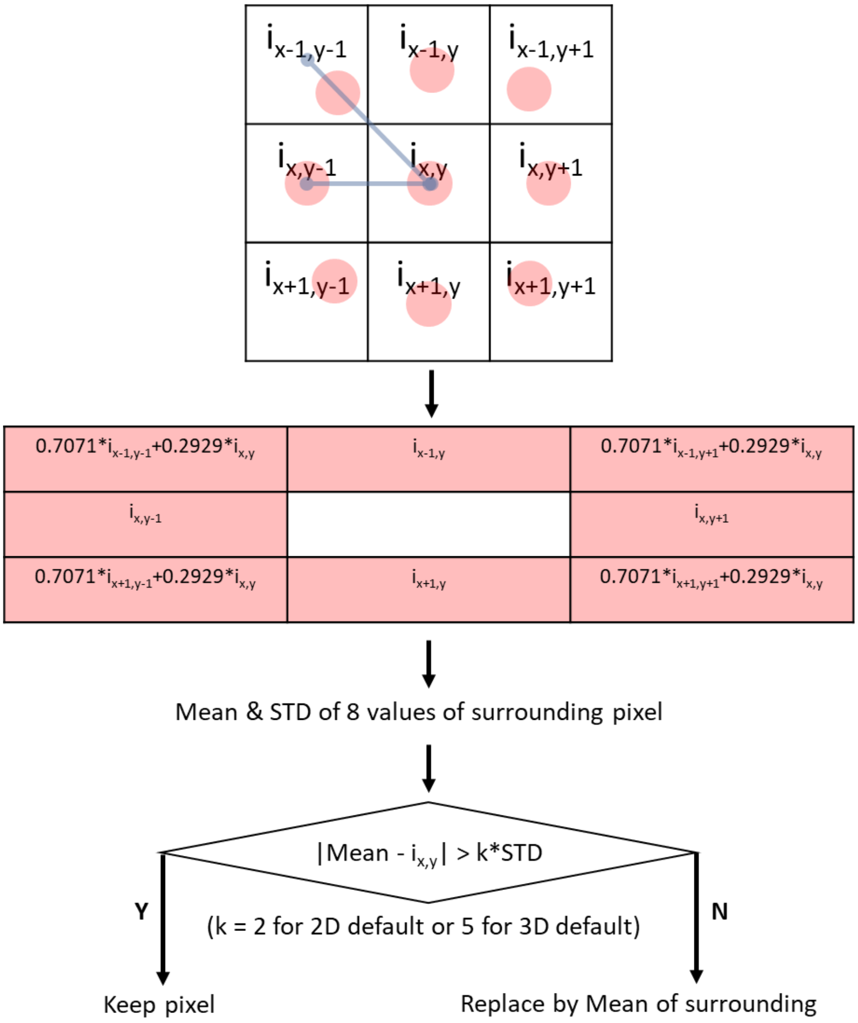


**Supplementary Figure 4. Significant difference filter.** Significant difference filter changes a pixel to the mean of its surrounding pixel if the object pixel is "significantly different" from its surrounding pixel. "Significant difference" is defined as a difference greater than 2-fold of standard deviation (STD) of surrounding pixels for 2D mode and 5-fold for 3D mode. The rationale behind is that there is no detectable independent actin element as tiny as one pixel element, so a one-pixel region that is significantly different from its surrounding tends to be noise or bias due to z-axis projection.

**Supplemental Figure 5: The mean and STD gradient magnitude of Ground Noise is directly proportional to STD of the Noise**

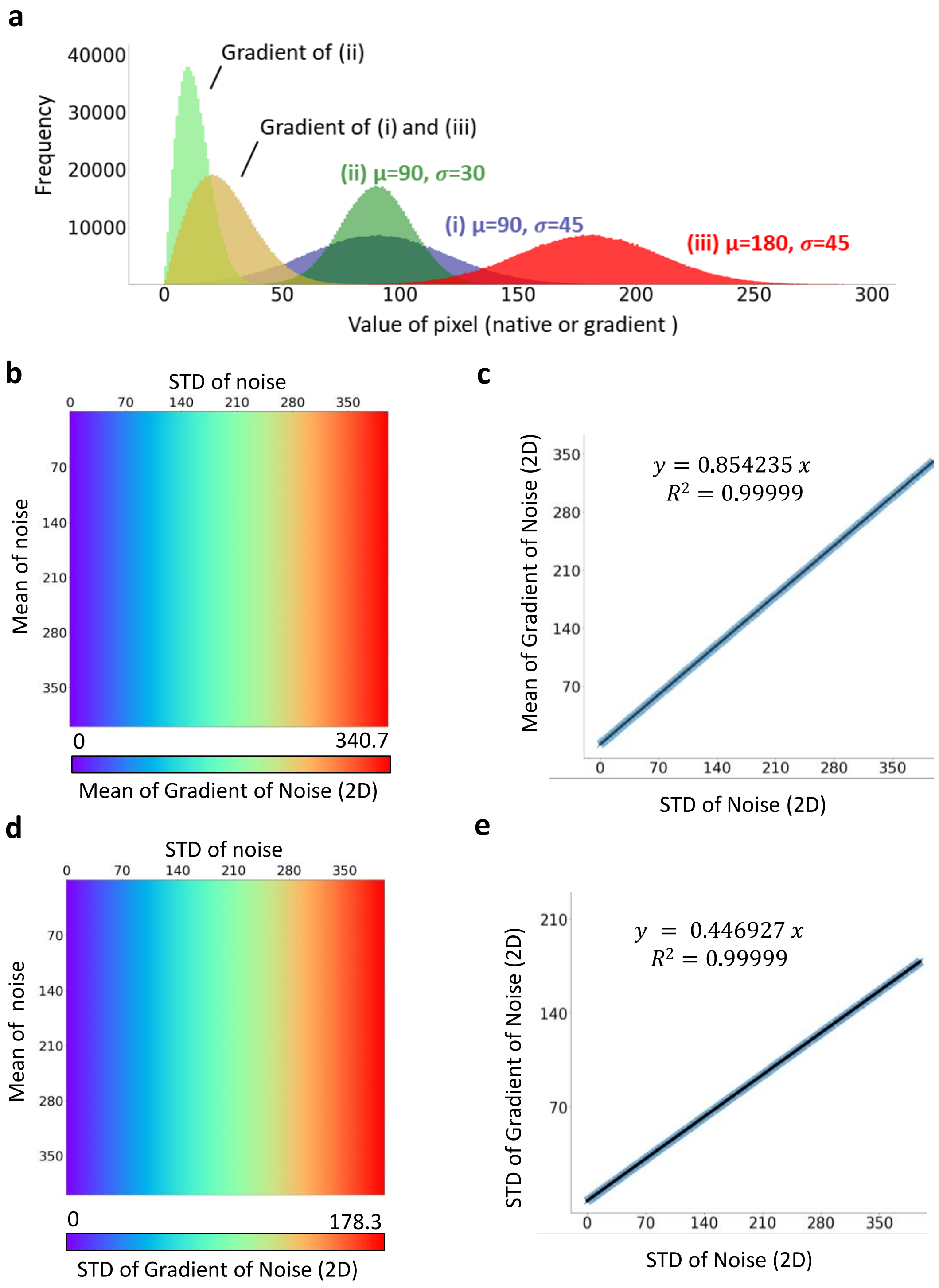


(see next page for legend)

**Supplementary Figure 5. The mean and standard deviation (STD) of gradient magnitude of ground noise is directly proportional to the STD of noise**. To investigate the mathematical relationship between statistical distribution of ground noise (represented by coarse background located by NNES) and the gradient magnitude of ground noise, we used random normal array that mimics the data structure of our image sample, and generated its Scharr gradient image for statistical analysis. **a,** Normal-like distribution of Scharr gradient**.** Simulated 800*800 ground noise image subject to normal distribution (i), (ii), and (iii) are processed by Scharr operator and the gradient distribution is shown. The gradient distribution is not influenced by the mean yet by the STD of native data of the simulated ground noise. **b** and **c**, The mean of noise gradient is not influenced by the mean of the noise, but the STD of noise. The mean of noise gradient is directly proportional to the STD of native noise. **d** and **e**, The STD of the noise gradient is only influenced by STD of the noise as well. The STD of the noise gradient is directly proportional to the STD of native noise.

**Supplemental Figure 6: The ratio of x-y unit and z unit influences the proportional coefficient of $\boldsymbol{\sigma_{Noise}}$-$\boldsymbol{\mu_G}$ and $\boldsymbol{\sigma_{Noise}}$- $\boldsymbol{\sigma_G}$ relationship.**

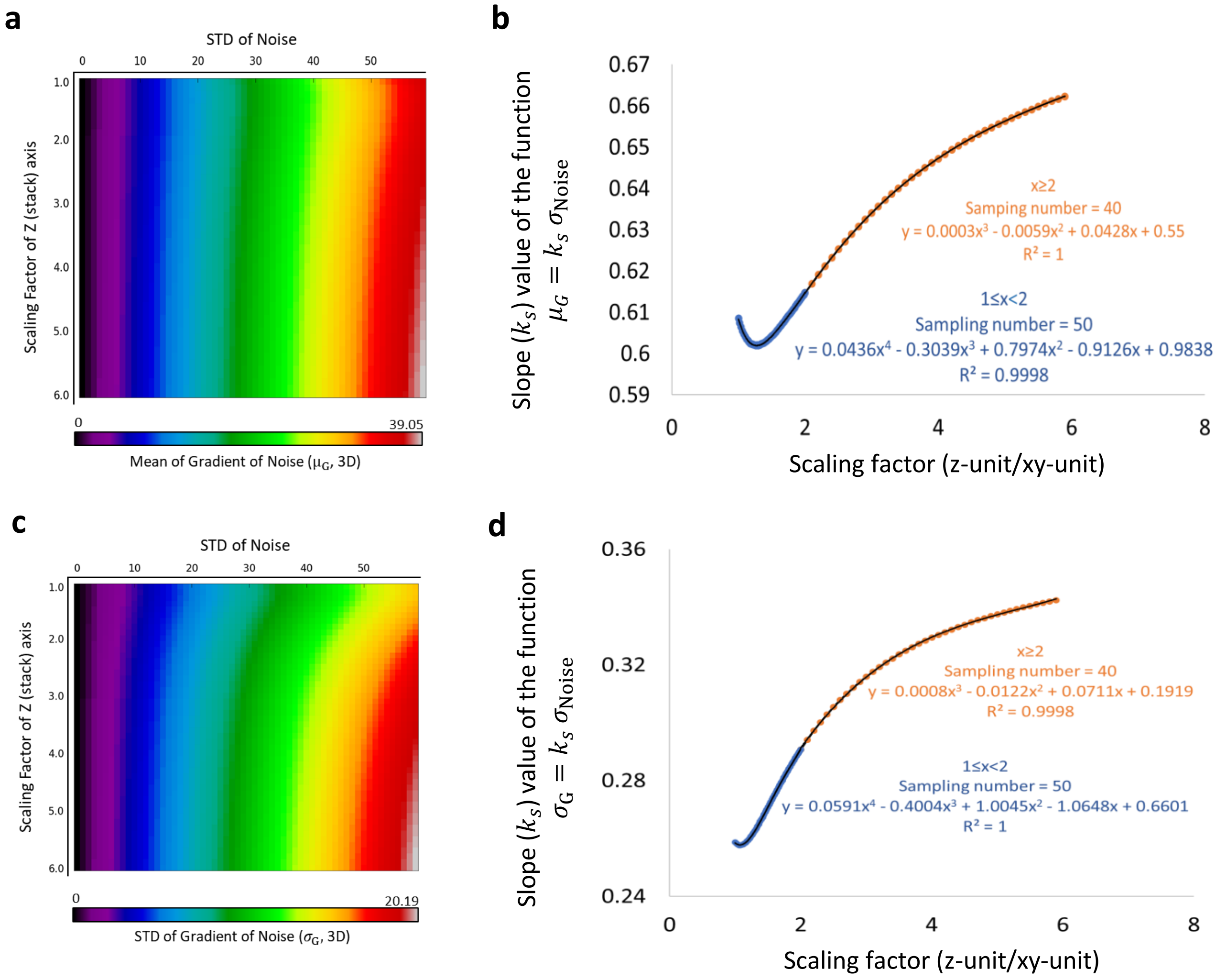


**Supplementary Figure 6. The ratio of x-y unit and z unit influences the proportional coefficient of $\boldsymbol{\sigma_{Noise}}$-$\boldsymbol{\mu_G}$ and $\boldsymbol{\sigma_{Noise}}$- $\boldsymbol{\sigma_G}$ relationship.** Using a strategy similar to what is presented in Supplementary Figure 5, we simulated 3D normal ground noise images (800*800*25) with a variable standard deviation (STD) and mean fixed to 90. Next, the z-axis was interpolated by the fold of a scaling factor, defined as z-unit / xy-unit, the second variable. Like the 2D data structure, at a given scaling factor, the mean and STD of noise gradient is proportional only to the STD of native noise. Then, we adopt a mathematical model $\mu_G$ or $\sigma_G = k_s(Scaling\ factor)\ \sigma_{Noise}$ to accurately describe the numeric relationship between the gradient and native value of background noise. By a piecewise polynomial regression, we obtain a prediction model that accurately ($R^2 = 1$) calculate $k_s$ and therefore determine the relationship between mean/STD of the noise gradient and the native noise values. **a** and **b**, the relationship between mean of noise gradient and STD of noise. **c** and **d**, the relationship between mean of noise gradient and STD of noise.

### Supplemental Figure 7: determination of global gradient threshold

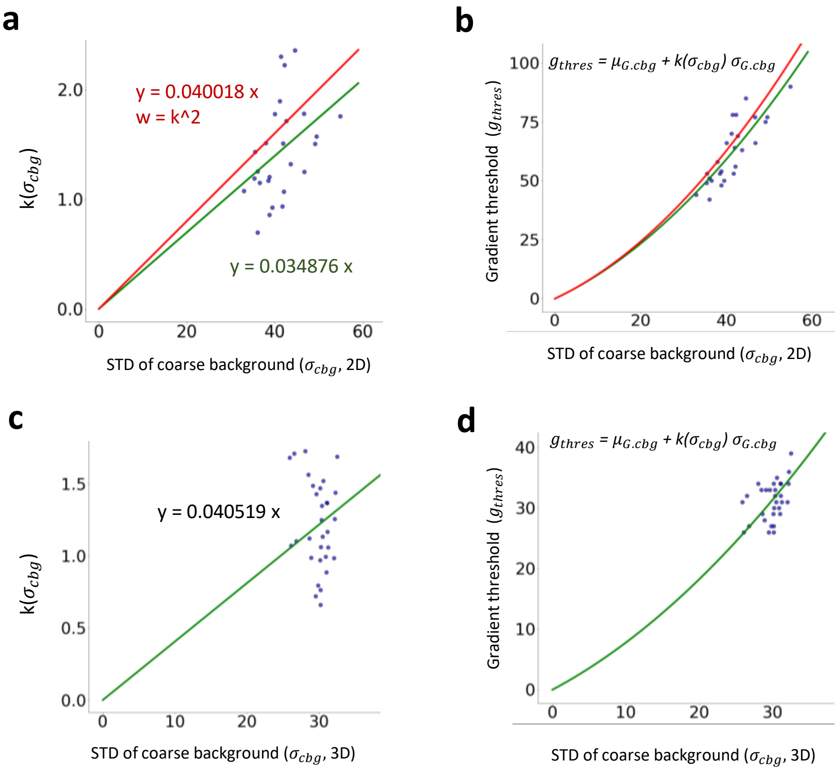


**Supplementary Figure 7. Determination of global gradient threshold.** In order to determine the global gradient threshold using estimated mean and standard deviation (STD) of coarse background gradient as an input of ILEE, we constructed a non-linear estimation model $g_{thres}$ = $\mu_{G.cbg}$ + $k(\sigma_{cbg})$ $\sigma_{G.cbg}$. The global gradient threshold of 30 samples with 800*800*25 resolution randomly collected from our actin image database were evaluated manually (similar to MGT), and the mean and STD of gradient values in coarse background are calculated following the approach described in the Methods and Supplementary Figure 5. For the 2D mode, images were first projected to 800*800 resolution. Then, the coefficient k of each sample was calculated and corelated with STD of the coarse background. **a,** The correlation of STD of coarse background and k. Green, standard mode, where each sample have the similar weight of regression; Red, conservative mode (default of this paper), where a weight equal to $k^2$ is applied. **b,** The $g_{thres}$-$\sigma_{cbg}$ relationship restored from a. **c** and **d,** Similar to **a** and **b**, but the reference global gradient threshold were manually evaluated using a 3D visualizing interface.

### Supplemental figure 8: impact of $K$ and the training for $K_2$ estimation

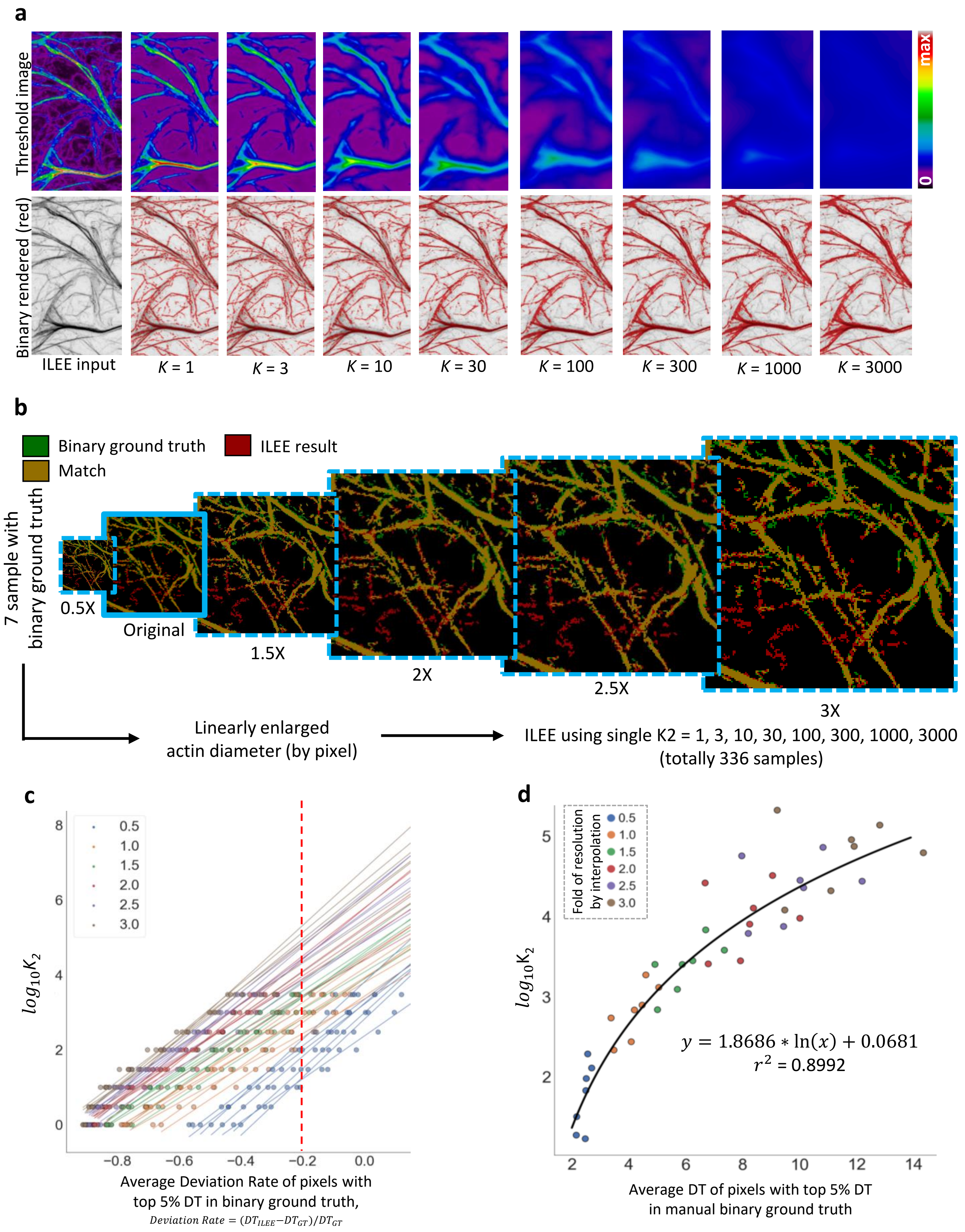


(see next page for legend)

**Supplementary Figure 8. The impact of *K* and the training for $K_2$ estimation.** In order to learn the impact *K* (implicit Laplacian smoothing coefficient) and determine the *K* value for a give batch of image sample, we constructed a training database and developed a non-linear estimation model. **a**, The influence of different *K* to rendered threshold image and binary result by ILEE. A low *K* renders a threshold image that assimilates the object sample with finer filament structure (preserved high frequency information), but tends to underestimate the thickness of bright and thick filament; as *K* increases, less local detail are preserved, and the rendered binary image losses thin and faint filaments but get more accurate for thick and bright filament. There is a trade-off over the performance over faint/thin and bright/thick for a single *K*. Therefore, we used the full-outer-join result of a small $K_1$ and a high $K_2$ (Fig. 3b and corresponding main text). **b**, Construction of $K_2$ training database. Initially, we planned using the 7 images with hand-portrayed binary ground truth of filament fraction, but these sample have relatively limited range of filament size by pixel, so we decided to expand out sample pools using these data. Each samples are bicubically interpolated into the resolution of 0.5-, 1,5-, 2-, 2.5-, and 3-fold of the original and added to the database. Their corresponding binary imaged are converted to float data type with 0.0 and 1.0 to process the same bicubic interpolation, with the pixels over 0.42 are defined as True and False if not. Using this approach, the judgement of matching between the ground truth and ILEE result did not have significant change, as shown. Finally, each sample will be processed by ILEE with a single $K_2$ ranging from 1 to 3000, rendering totally 336 binary images as the training database. **c** and **d**, Training algorithm. In **c**, we converted the ILEE samples to a feature value – estimated $K_2$ with -0.2 of average deviation rate of pixels with top 5% DT in binary ground truth. Specifically, for each original or interpolated image sample (shown by independent lines, where different color represents different fold of interpolation), ILEE binary results using various single $K_2$ are compared with corresponding ground truth image to calculated the deviation rate, defined as the fold of difference of Euclidian distance transformation (DT) of ILEE result vs ground truth relative to ground truth. The averaged deviation rate of pixels with top 5% DT are taken as a feature value to represent thick filament. Next this feature was correlated to the corresponding $K_2$, and linear regressions were conducted to estimate the $\log_{10}$ of the optimal $K_2$ that renders -0.2 for average deviation rate for each sample. Finally, the estimated optimal $K_2$ and the average DT of pixels with top 5% DT of each sample were utilized (shown in **d**) to generated an exponential regression model. This model covers the vast majority (if not all) of possible highest filament thickness rendered by confocal microscope images of cytoskeleton. The average DT of pixels with top 5% DT estimated by Niblack thresholding were used as an independent variable for optimal $K_2$ estimation.

#### Supplementary Figure 9: Visualized demonstration of concepts of cytoskeleton indices.

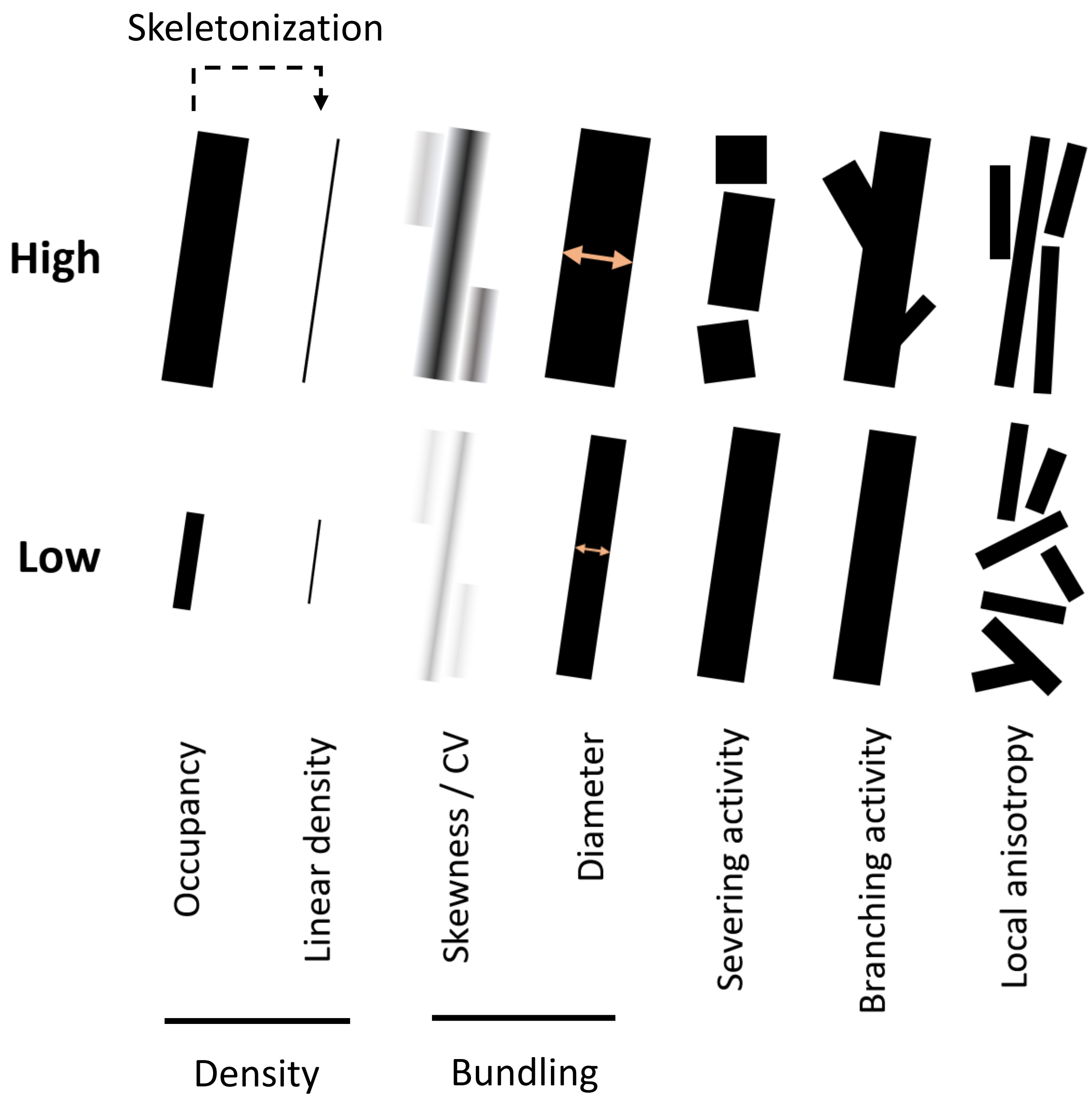


**Supplementary Figure 9. Visualized demonstration of concepts of cytoskeleton indices.** Major cytoskeleton indices involved in this research are explained by showing schematic cytoskeleton pieces representing high vs low value of the indices. Skewness/CV, as well as Diameter(TDT/SDT) have similar concept and reflect similar features of cytoskeleton image, with minor difference in their mathematical definition (see ***Methods***); therefore, their demonstrations are merged.

**Supplemental Figure 10: The stability of ILEE and other classic image thresholding approaches for cytoskeleton segregation in confocal images.**

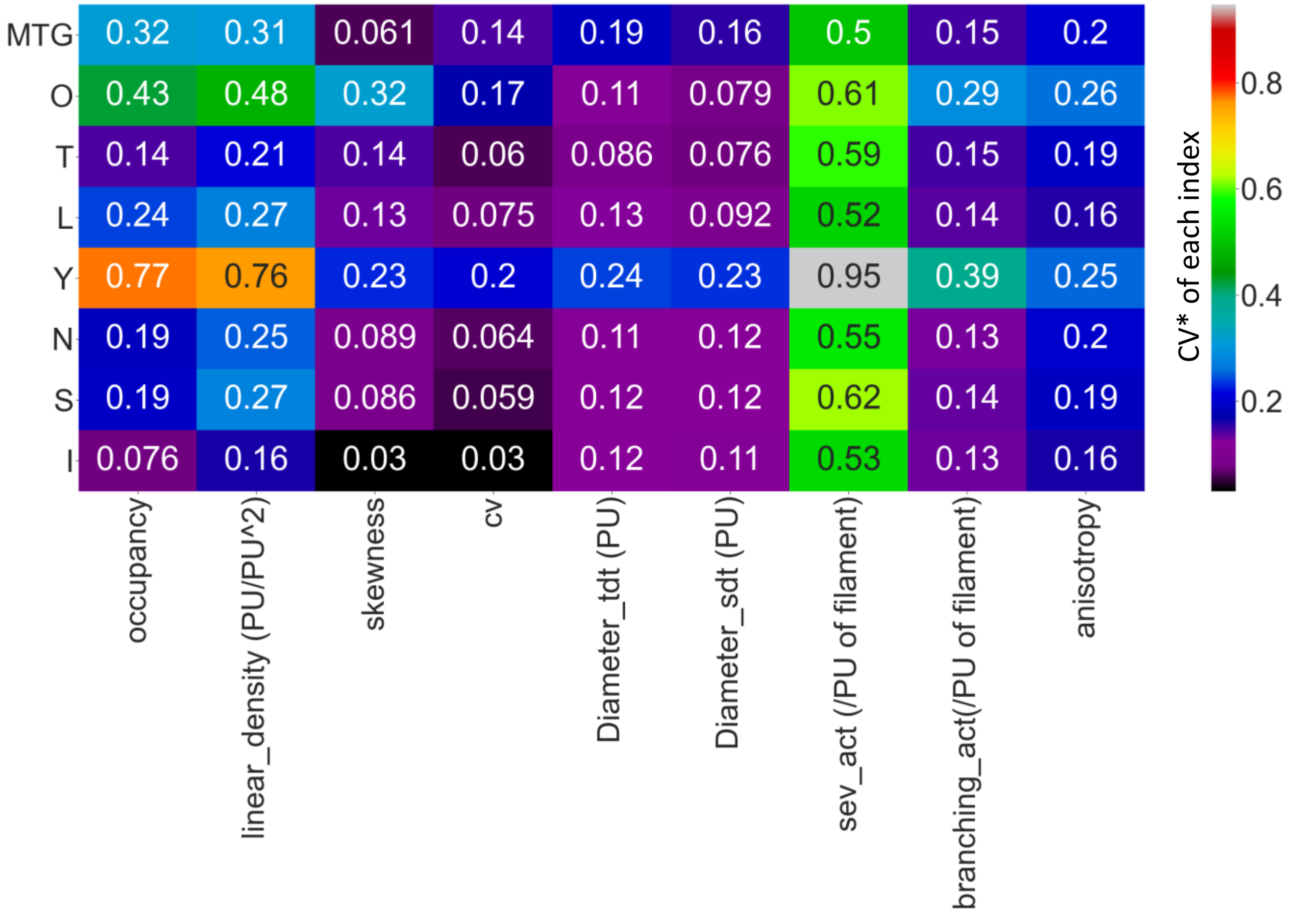


**Supplementary Figure 10. The stability of ILEE and other classic image thresholding approaches for cytoskeleton segregation in confocal images.** The coefficient of variation (CV*) of each index rendered by each approach in Fig. 4**d** were calculated and visualized as a heatmap. O, Ostu; T, Triangle; L, Li; Y, Yan; N, Niblack; S, Sauvola; I, ILEE. ILEE has a dominantly low CV* (i.e., high stability) over diverse sample for *occupancy*, *linear density*, *skewness*, *CV*, or one of the lowest for other indices. Note that the CV* of computed indices in the figure and the fluorescence *CV* of a sample, which is a cytoskeletal index, are different concept.

**Supplemental Figure 11: The visualized comparison of robustness of ILEE and other algorithms by segmentation accuracy.**

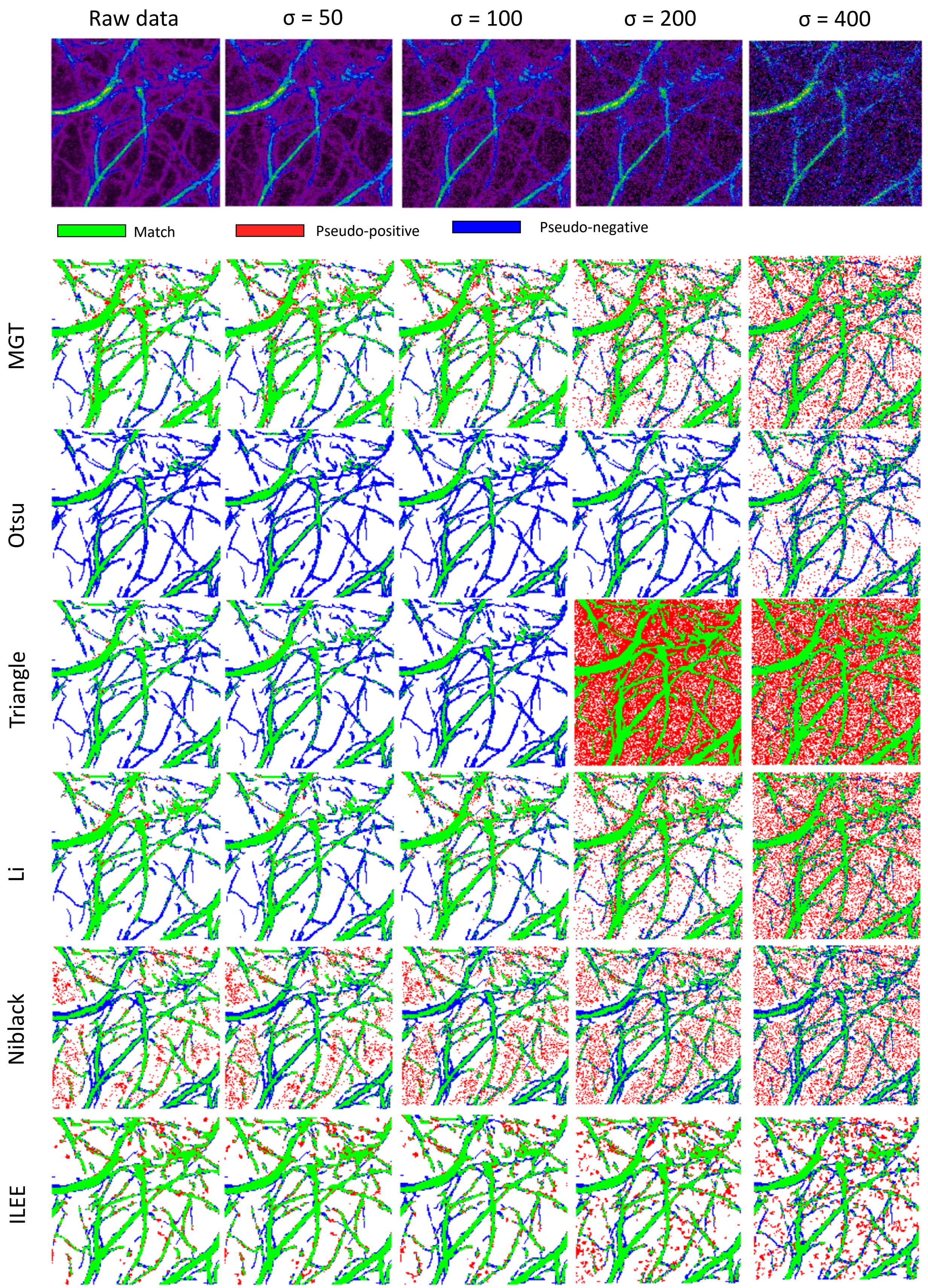

**Supplementary Figure 11. The visualized comparison of robustness of ILEE and other algorithms by segmentation accuracy.** The presentation image **Figure 4-a** are added with a series of gaussian noise ($\mu$ = 0, $\sigma$ as variable). The binary images are computed by selected algorithms including ILEE and compared with manually portraited ground truth, and pixels of match, pseudo-positive, and pseudo-negative are presented. By visual observation, all of the algorithms are stable when the $\sigma$ of noise is within 100 (3~4% of max of dynamic range); 3 of the better performed algorithm, ILEE, MGT, and Li, remain accurate, among which ILEE has the best coverage of ground truth. When $\sigma$ of noise is higher than 100, all algorithms becomes visibly unstable. While MGT and Li tend to have single pixel errors while ILEE tends to result in block-shaped errors that are less in number and mimics the thickness of the cytoskeleton in shape, which indicates that the indices derived from ILEE are potentially more accurate at high noise conditions.

### Supplemental Figure 12: The quantificational comparison of robustness of ILEE and other algorithms by segmentation accuracy.

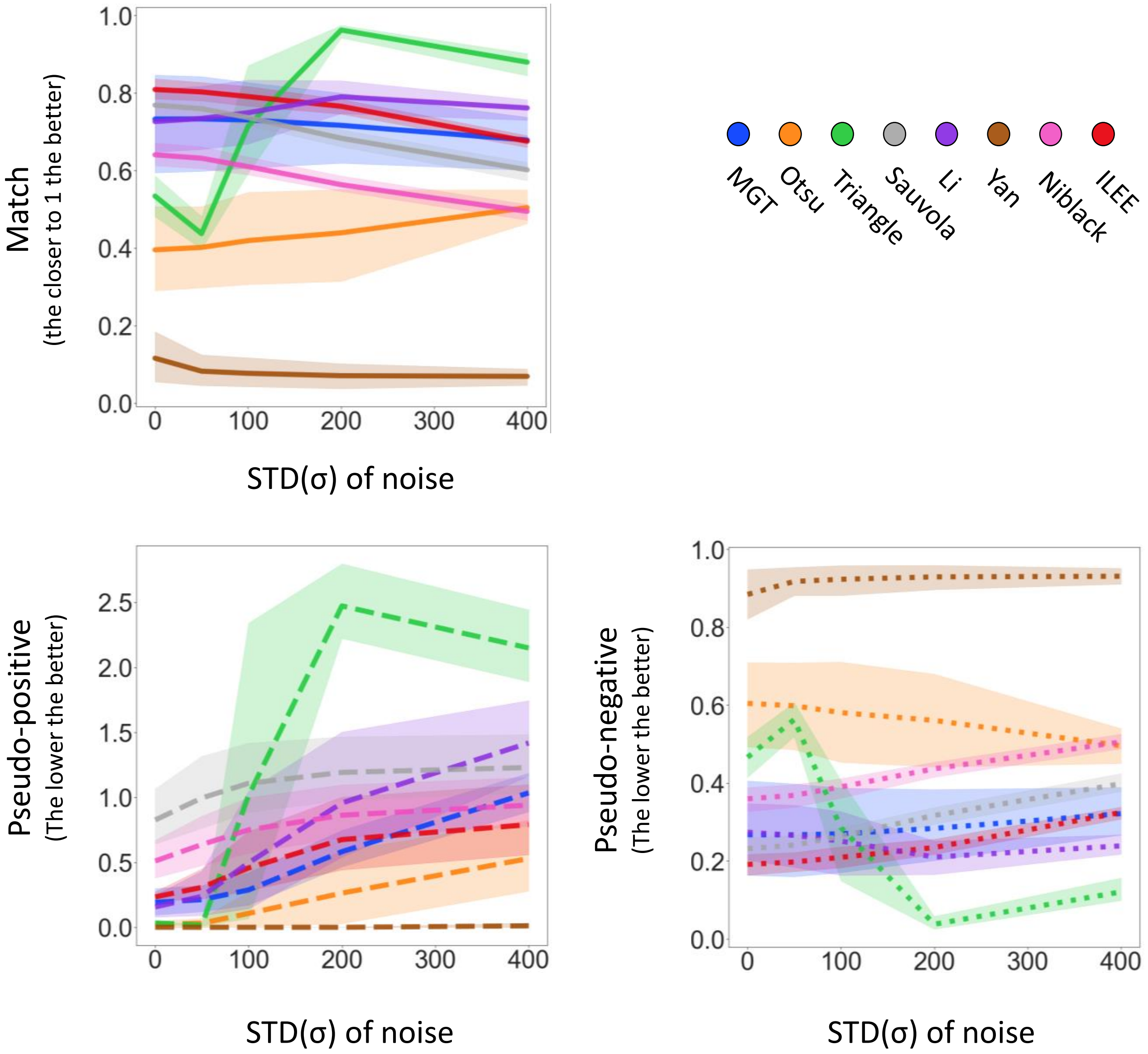


**Supplemental Figure 12: The quantificational comparison of robustness of ILEE and other algorithms by segmentation accuracy.** The dataset of **Figure 4-b,c,d** are added with a series of gaussian noise (μ = 0, σ as variable). Since gaussian noise is random and potential unstable, each sample-noise combination are technically repeated by 12 times and the averaged result is used. The transparent area with light color indicates 95% confidence interval of each algorithm. The binary images rendered by different algorithms are compared with manually portraited ground truth to count the pixels that are matched (ideally 1.0), pseudo-positive (ideally 0), and pseudo-negative (ideally 0). Generally speaking, ILEE is the best-performing algorithm with high match, low pseudo-positive, and low pseudo-negative rate at both low and high noise. Other algorithms have one or more flaws.

**Supplemental Figure 13: The quantificational comparison of robustness of ILEE and other algorithms by index rendering stability.**

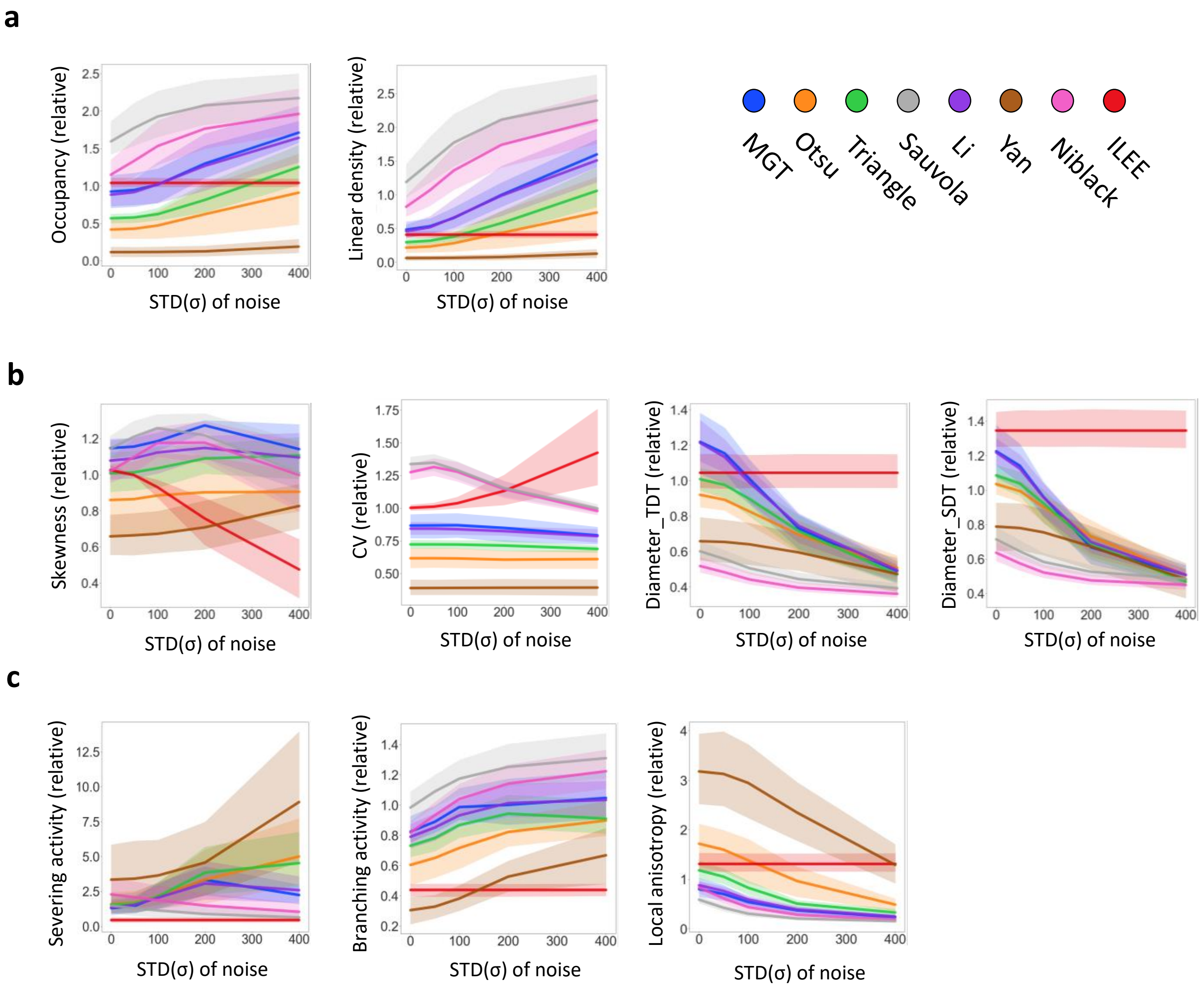


**Supplemental Figure 13: The quantificational comparison of robustness of ILEE and other algorithms by index rendering stability.** Like Supplemental Figure 12, the dataset of **Figure 4-b,c,d** are added with a series of gaussian noise (μ = 0, σ as variable), segmented by aforementioned algorithms, and their indices are computed and presented as the relative value to the ground truth. Each sample-noise combination are technically repeated by 12 times and the averaged result was used. The transparent area with light color indicates 95% confidence interval of each algorithm. For each figure, the lines with plainer slope indicate higher robustness (resistance to noise); being closer to 1.0 by value indicates higher accuracy. **a**, Index class of density; **b**, index class of bundling; **c**, indices of other classes. For most of the indices, ILEE provide an extremely stable result against increasing noise, while other algorithms has very obvious change of value of output indices and are therefore no longer accurate (if they were), which echoes the visual observation of **Supplemental Figure 12**. Interestingly, *skewness* and *CV* are two exceptions, where ILEE shows more instability and tends to have a bifurcated direction of change.

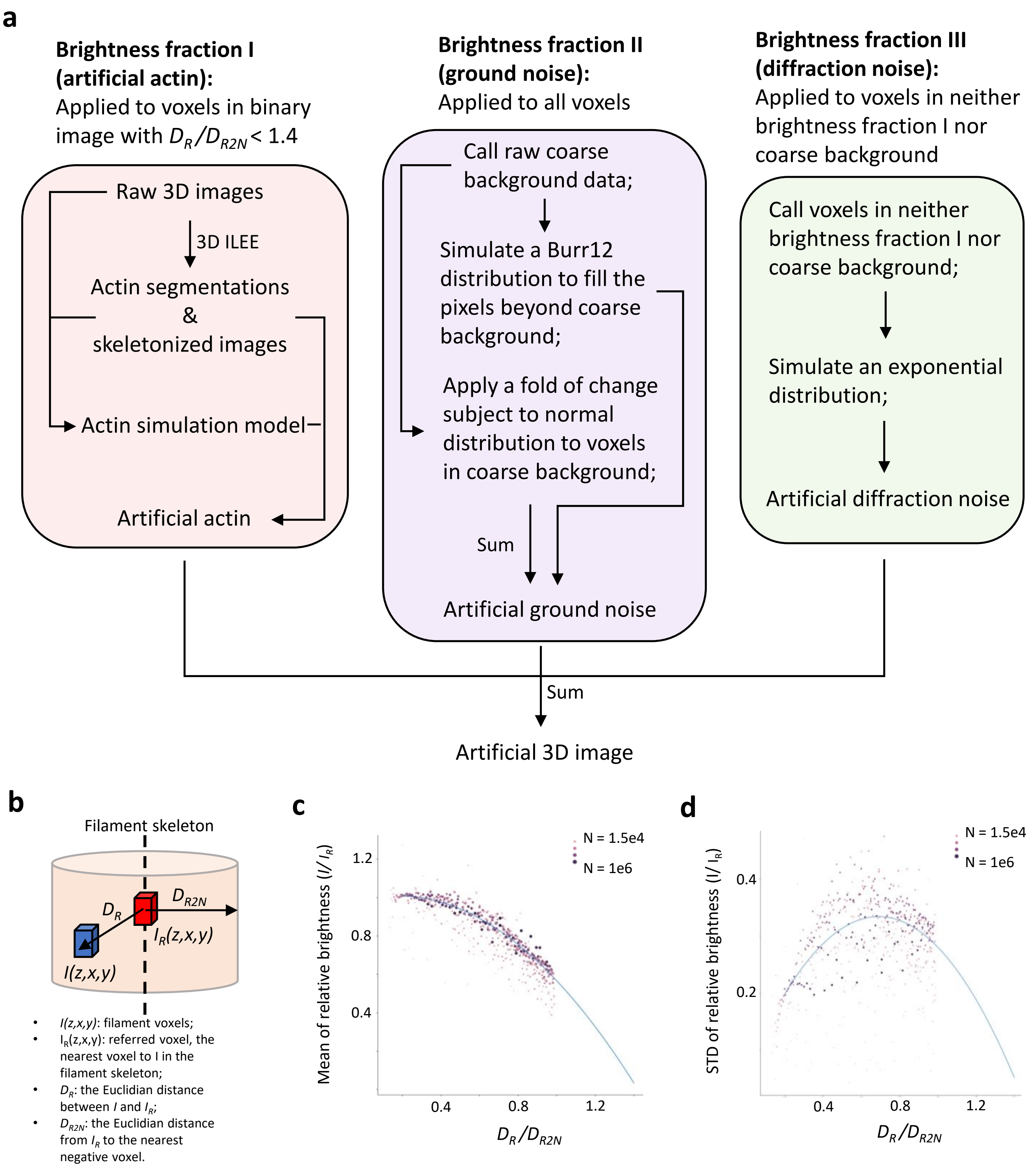


(see next page for legend)

**Supplemental Figure 14: an illustrated introduction of the actin image simulation model to generate 3D cytoskeleton images with ground truth. a**, a schematic diagram describing the image simulation model. The training dataset is the same as Fig. 2. As preparation, raw images were processed by 3D ILEE. To mostly simulate the shape of actin, we directly adapted the skeletonized images ($I_{sk}$) generated from 3D ILEE as the central line of new filaments but filled all voxels with novel values. The artificial images comprise three brightness fractions: artificial actin, artificial ground noise, and artificial diffraction noise. First, referring to the binary images of the training samples, for each voxel ($I$) in the filament, we found its referred voxel ($I_R$), the nearest voxel to $I$ in the filament skeleton. Next, we calculated the $D_R$ (the Euclidian distance between $I$ and $I_R$) and $D_{R2N}$ (the Euclidian distance from IR to the nearest negative voxel) of all voxels that belongs to the filament (also see **b**) and constructed a polynomial regression model (also see **c**) between the relative distance to $I_R$ (i.e., $D_R / D_{R2N}$) and the mean ($\mu_{atf}$) as well as the STD($\sigma_{atf}$) of relative brightness (i.e., $I / I_R$) for all on-filament voxels for all images. To generate the new artificial actin fraction, we refilled the on-filament voxel by the normal distribution $I / IR \sim N(\mu_{atf}, \sigma_{atf}^2)$ based on the brightness of voxels belonging to the corresponding $I_{sk}$, and we abandoned the voxels with $D_R / D_{R2N}$ >1.4 because they are more likely to be the structural noise – the cytosol and PM with fluorescence markers not binding to the actin. Second, for the ground noise fraction, we had different strategies to generate two groups of voxels. For the volume previously belonging to the coarse background, we directly apply a fold of change that is subject to a normal distribution to each voxel; for the other volume beyond coarse background, we gave random values that subject to a Burr-12 distribution fitted by voxels of the coarse background, thereby all voxels in the new artificial image were given with a ground noise value. Third, the volume previous belonging to neither coarse background nor real actin is considered as the space impacted by the diffraction (Fig. 2, **a**). Using the raw data of this region, we fitted an exponential distribution to describe the diffraction signal and refilled the diffraction region by this distribution. The choice of Burr-12 and exponential distribution is based on the best performance over a fitting test over 60 statistical models. Finally, we add the three fractions of brightness together and linearly align the value range to 0-4095 (12 bit). **b**, a schematic diagram explaining the data utilized to fit the actin simulation model. **c**, **d**, the fitting result of the expected Mean of STD of the relative brightness of the actin simulation model. Only data with $D_R / D_{R2N}$ lower than 1 were used to fit the model because they were less likely to be influenced by the structural noise. As $D_R / D_{R2N}$ were discretely distributed, we added weights equal to $\log_{10}N$, where N means the number of voxels among the 31 samples, to each $D_R / D_{R2N}$.

**Supplemental Figure 15: The stability/robustness of ILEE and MGT in batch analysis of biological samples of Figure 5**

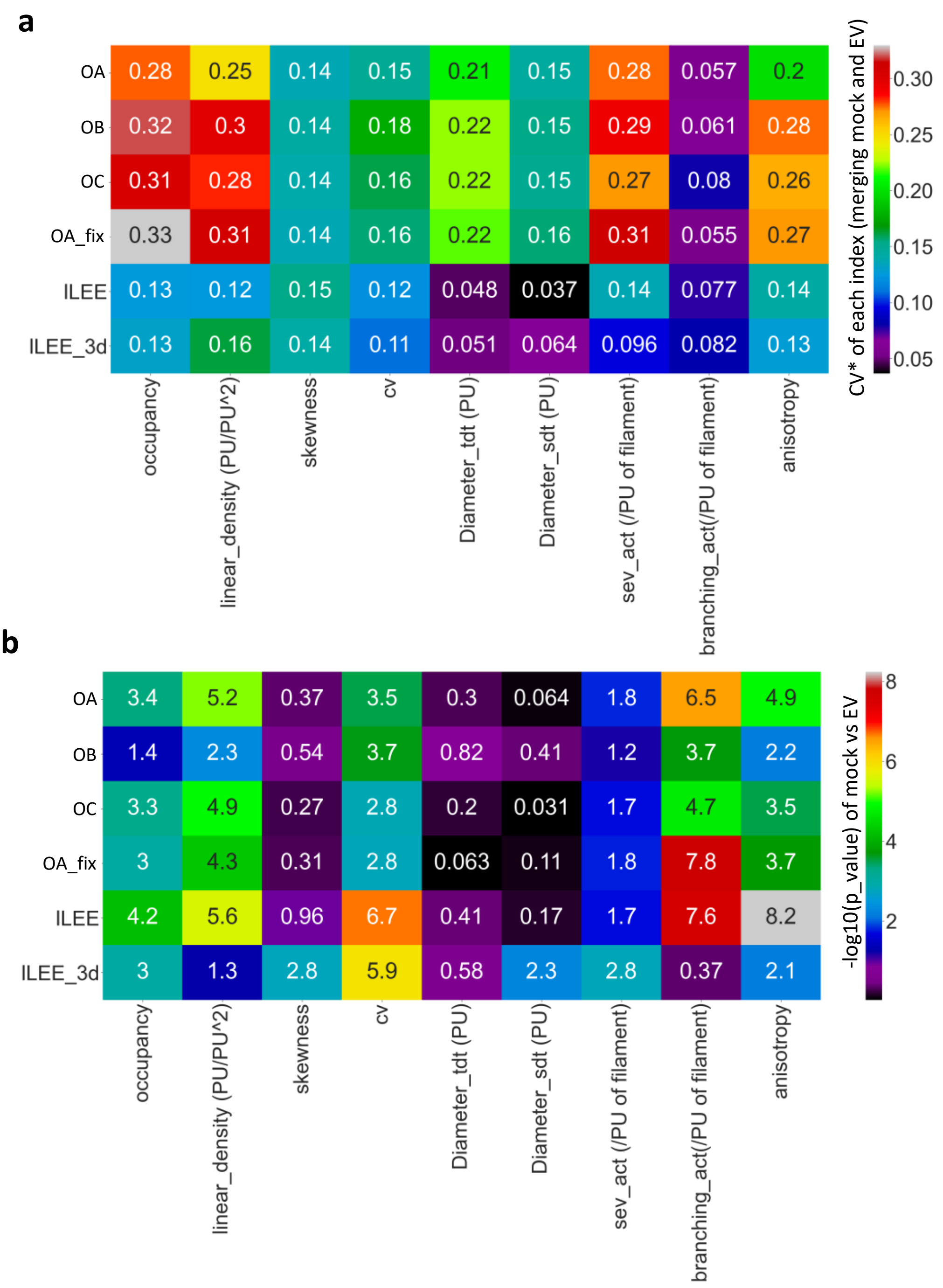


**Supplementary Figure 15. The stability/robustness of ILEE and MGT in batch analysis of biological samples of Figure 5. a**, The coefficient of variation (CV*) of each index rendered by ILEE and MGT by different operators. The individual CV*s of mock and EV group for each index-method combination are merged. ILEE or ILEE_3d got the lowest CV* (or highest stability) on *occupancy*, *linear density*, (fluorescence) *CV*, *Diameter_TDT/SDT*, *static severing activity*, and *local anisotropy*. **b**, A comparison of the t-test P value of mock versus *P. syringae* (EV)-inoculated sample rendered by ILEE and MGT by different operators. ILEE or ILEE_3d has the highest $-\log_{10}$ P value (i.e., lowest P value) on *occupancy*, *linear density*, *skewness*, (fluorescence) *CV*, *Diameter_SDT*, *static severing activity*, and *anisotropy*. Note that the CV* of computed indices in the figure and the fluorescence *CV* of the samples, which is a cytoskeletal index, are different concept.

**Supplemental Figure 16: the correlation of *occupancy* and *linear density***

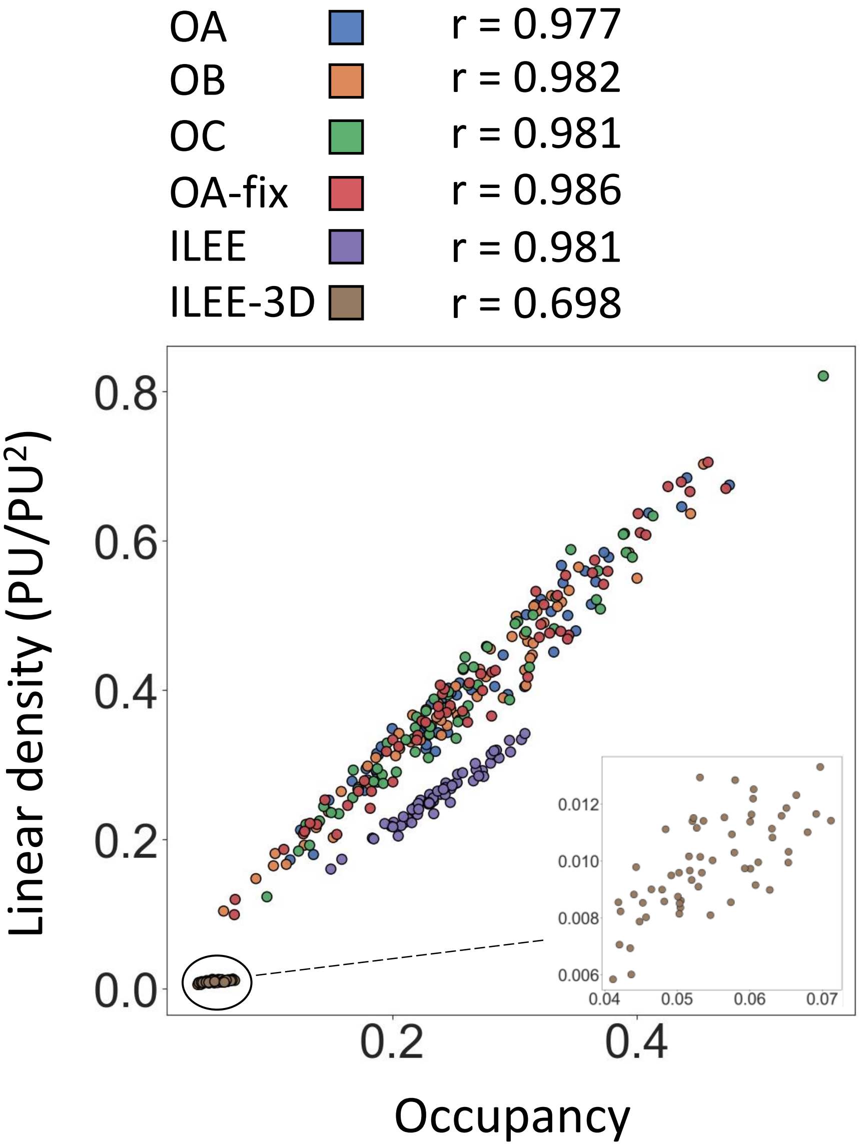


**Supplementary Figure 16. The correlation of *occupancy* and *linear density*.** The *occupancy* and *linear density* rendered by ILEE and MGT shown in Figure 5 are corelated. The *occupancy* and *linear density* generally have a very high linear correlation, indicating that they agree with each other on evaluating the cytoskeletal density. ILEE and MGT does not display the same mathematical relationship, indicating that ILEE and MGT possess different tendencies in rendering the topological structure (mostly influencing linear density). In the 3D mode, they only have medium-strong correlation, potentially because the 3D topological structure cannot be perfectly rendered due to concave/convex structures present in the skeletonized images. Additionally, this could be the result of abandoned oversampling process for generating the skeletonized image due to insufficient computational power for standard PCs.

#### Supplemental Figure 17: The performance of ILEE on other types of biological sample

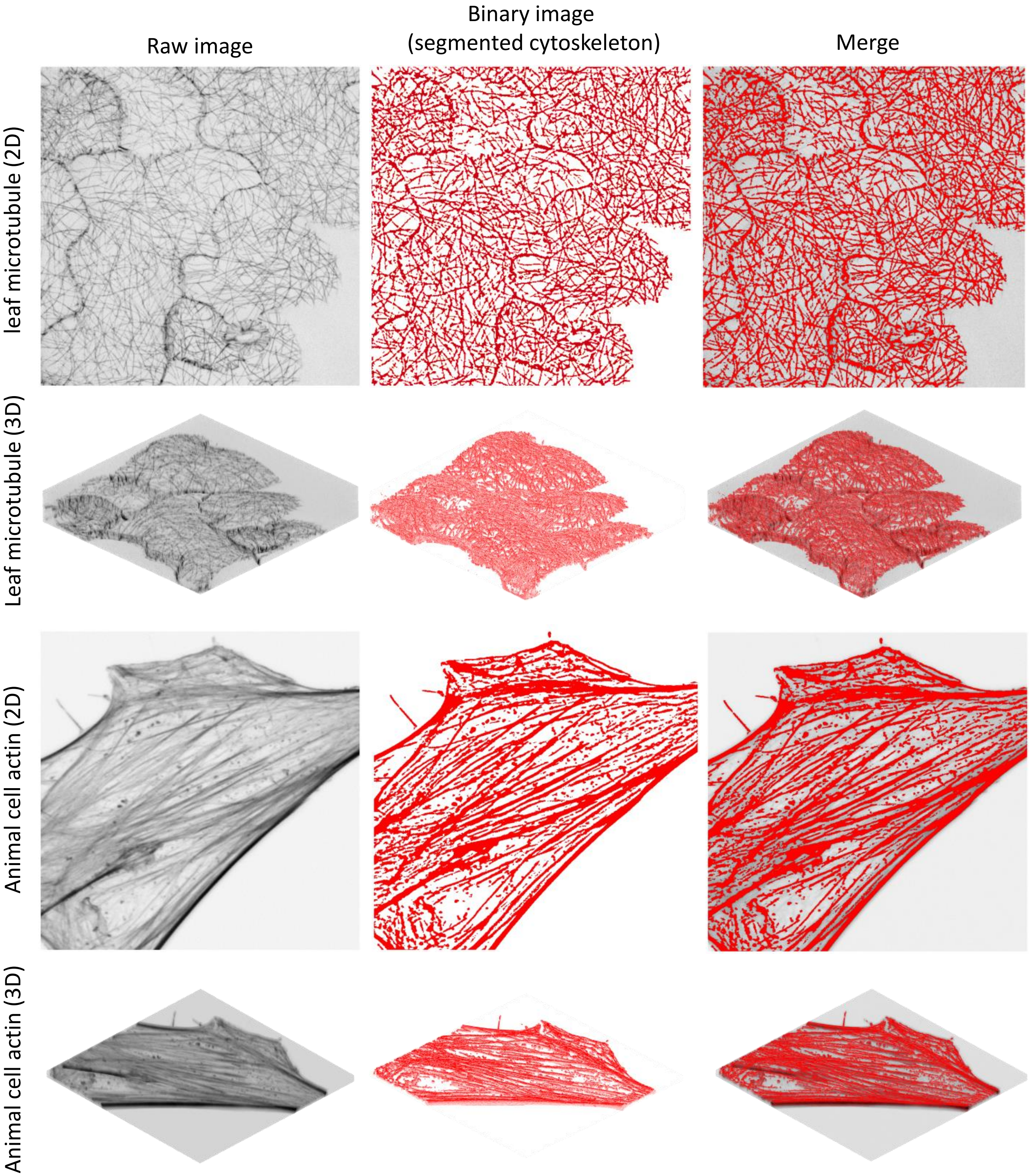


**Supplementary Figure 17. The performance of ILEE on other type of biological sample.** We tested ILEE on leaf microtubule images of Arabidopsis Col-0/GFP-MAP4 and animal cell actin images of YUMMER1.7D4 line. The Col-0/GFP-MAP4 images were generously provided by Dr. Silke Robatzek (Ludwig-Maximilians-Universität München) from a previously published dataset[35]. The result indicates ILEE accurately covers the cytoskeleton samples with locally dynamic brightness, thickness, and shape, with no visible difference compared with its performance on leaf actin images.

**Supplemental Figure 18: ILEE and human eye have different tendency to judge the topological structure of cytoskeleton, especially between two bright bundles**

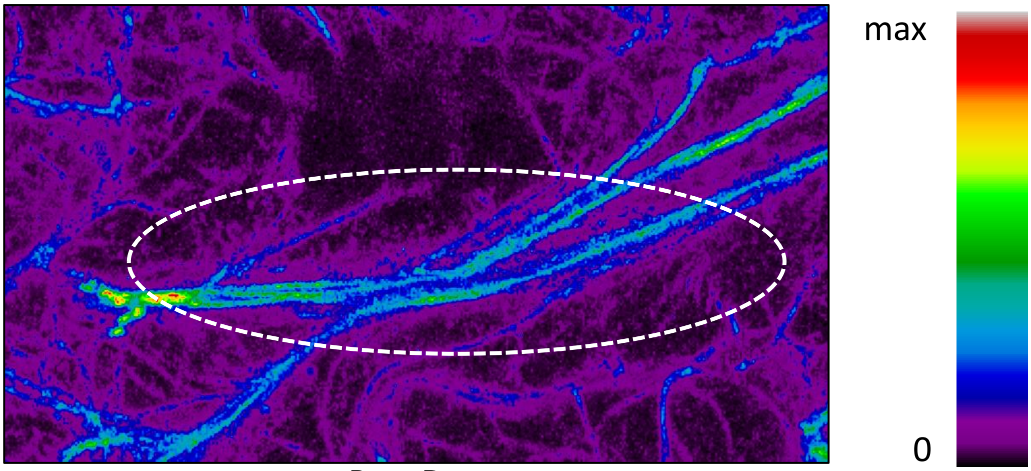


Raw Data

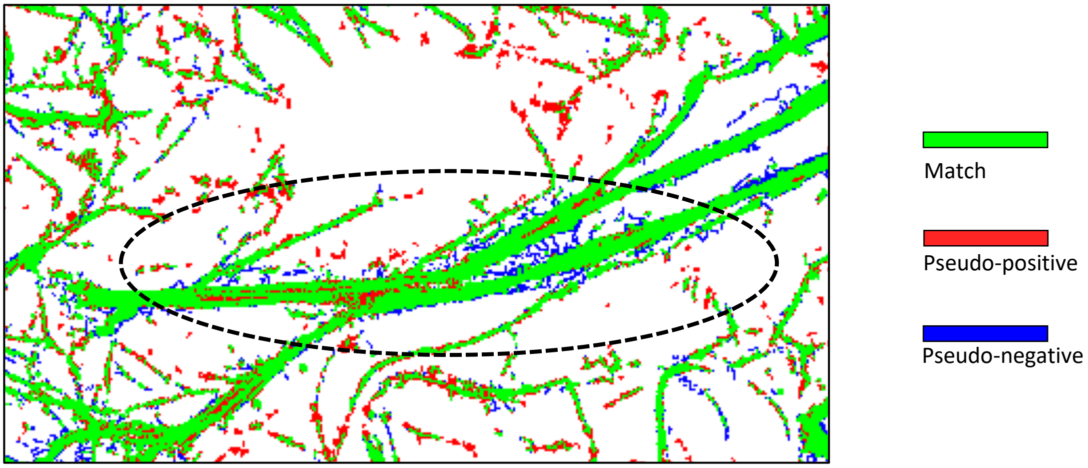


ILEE vs ground truth comparison of binary image

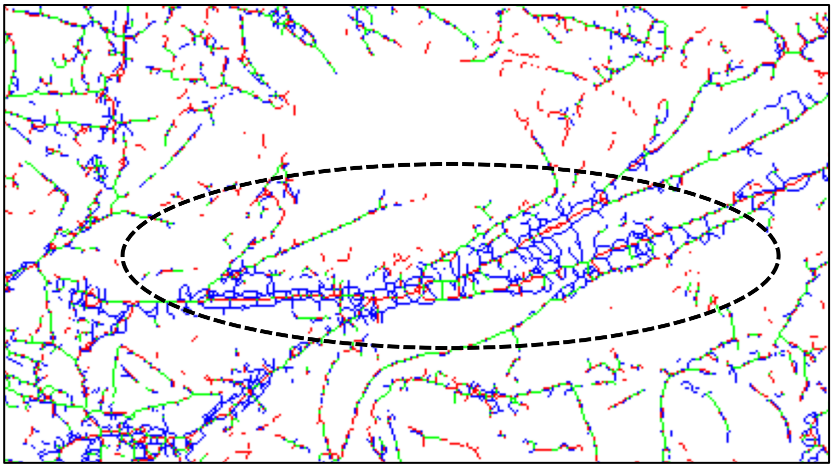

ILEE vs ground truth comparison of skeleton image

(see next page for legend)

**Supplementary Figure 18. ILEE and human eye have different tendency to judge the topological structure of cytoskeleton, especially between two bright bundles.** A cropped portion of the demonstration image from Fig. 3**a** and its derived images are generated and demonstrated to explain the discrepancy related to the determination of the topological structure between ILEE and hand-drawn binary ground truth (i.e., human eye evaluation). When ILEE binary image is compared with ground truth, they mostly match with each other with only minor and non-influential differences. When both binary images were skeletonized based on their topological structure, a numerous branches were detected from the ground truth, but not from ILEE. This phenomenon is very obvious in the highlighted (i.e., circled) area, where two bright actin bundles are very close to each other. This is intriguing, as where the human eye identified many branches and intervals, ILEE did not. These structures have a strong impact on the topological structure. Therefore, the topology-sensitive indices, *linear density*, *severing activity*, and *branching activity* display contrast and stable inconformity between ground truth and ILEE (Fig. 4**d**). However, since the hand-drawn ground truth image is not a rigorously defined ground truth, we surmise that inconformity cannot be interpreted to any inaccuracy of ILEE.
